# Evolutionary conserved RLF, a plant cytochrome *b*_5_-like heme-binding protein, is essential for organ development in *Marchantia polymorpha*

**DOI:** 10.1101/2024.09.02.610766

**Authors:** Kentaro P. Iwata, Takayuki Shimizu, Yuuki Sakai, Tomoyuki Furuya, Hinatamaru Fukumura, Yuki Kondo, Tatsuru Masuda, Kimitsune Ishizaki, Hidehiro Fukaki

**Author notes:** Author for correspondence Hidehiro Fukaki.

## Abstract

In *Arabidopsis thaliana*, REDUCED LATERAL ROOT FORMATION (RLF), a cytochrome *b*_5_-like heme-binding domain (Cytb5-HBD) protein, is necessary for proper lateral root formation. Whereas the other Cytb5-HBD proteins in *A. thaliana* regulate different metabolic reactions, RLF is unique as it specifically regulates organ development. However, it remains unknown whether heme binding to RLF is necessary for its function, and whether *RLF* orthologs in different plant species also regulate organ development. We demonstrate that RLF binds to heme *in vitro* and that the two histidine residues, which are conserved among Cytb5-HBD, are crucial for both heme binding and its biological function in *A. thaliana*. In addition, Mp*RLF*, a *RLF* ortholog in *Marchantia polymorpha*, rescues the lateral root formation phenotype of the *A. thaliana rlf* mutant. Mp*rlf^ge^,* the loss-of-function mutation in the Mp*RLF*, resulted in delayed thallus growth and inhibited both gemma cup and antheridiophore formation. Transcriptome analysis using Mp*rlf^ge^* revealed that Mp*RLF* affects several metabolic pathways. Our findings indicate that Mp*RLF* is essential for vegetative and reproductive development in *M. polymorpha*, suggesting that the RLF-dependent redox reaction systems are evolutionarily conserved as crucial mechanisms for organ development across diverse plant species.

## Introduction

Heme is a porphyrin complex with a centrally coordinated iron atom. In plants, heme shares a metabolic pathway with chlorophyll biosynthesis up to the production of protoporphyrin IX, after which ferrochelatase coordinates Fe^2+^ to protoporphyrin IX (Roper & Smith, 1997; Hederstedt, 2012). In the Protein Data Bank (URL https://www.rcsb.org), which contains a database of 222,415 proteins, 4,272 protein chains have been identified as heme-binding proteins (HBPs) from entries containing heme type *b* and *c* (Li *et al*., 2011). The binding of these HBPs to heme is maintained by the coordination of specific amino acids, such as histidine, within the apoprotein to the iron atoms of the heme. In general, in various organisms, HBPs are essential for diverse biological processes such as steroid biosynthesis, aerobic respiration, and programmed cell death because of their role in electron transfer, substrate oxidation, and metal ion storage (Reedy & Gibney, 2004; Layer *et al*., 2010). In plants, HBPs such as cytochrome *c*, which is involved in electron transfer during photosynthesis, and SOUL4, which is implicated in lipid metabolism within chloroplast fat droplets (plastoglobule), have demonstrated diverse physiological effects (Kerfeld & Krogmann, 1998; Shanmugabalaji *et al*., 2020). Recently, proteomic analyses in *Arabidopsis thaliana* have identified a variety of HBPs, such as basic/helix-loop-helix type nuclear transcription factors and intracellular signaling factors involved in GTPase activation (Shimizu *et al*., 2020). These studies have revealed that HBPs function as intracellular signals regulating diverse physiological functions. However, it remains unclear how they are involved in plant organ development. One group of HBPs in plants is a family of proteins with a cytochrome *b*_5_-like heme-binding domain (Cytb5-HBD). In plants, the metabolic pathways mediated by most Cytb5-HBD proteins are diverse, including fatty acid desaturation, lignin biosynthesis, and nitrate reduction (Nagano *et al*., 2012; Gou *et al*., 2019). The genome of *A. thaliana* has 15 proteins with a Cytb5-HBD including five members of cytochrome *b*_5_ family [At1g26340 (CB5A), At2g32720 (CB5B), At2g46650 (CB5C), At5g48810 (CB5D) and At5g53560 (CB5E)] (Maggio *et al*., 2007), one cytochrome *b*_5_ like protein [At1g60660 (CB5LP)], four membrane-associated progesterone binding proteins [At2g24940 (MAPR2), At3g48890 (MAPR3), At4g14965 (MAPR4) and At5g52240 (MAPR5)] (Yang *et al*., 2005), two Δ-8 sphingolipid desaturases [At3g61580 (SLD1) and At2g46210 (SLD2)] (Sperling *et al*., 1998), two nitrate reductases [At1g77760 (NR1) and At1g37130 (NR2)] (Cheng *et al*., 1988; Wilkinson & Crawford, 1991) and At5g09680/REDUCED LATERAL ROOT FORMATION (RLF) (Ikeyama *et al*., 2010).

Whereas most of the Cytb5-HBD proteins in *A. thaliana* regulate a variety of metabolic reactions, At5g09680/RLF is an unique Cytb5-HBD protein that regulates organ development, including lateral root (LR) formation (Ikeyama *et al*., 2010). In general, LR formation is essential for root system architecture in most vascular plants and is regulated by auxin. In *A. thaliana*, LR formation is regulated by several auxin signaling modules including the SOLITARY-ROOT (SLR)/IAA14, an auxin/indole-3-acetic acid (Aux/IAA) protein - AUXIN RESPONSE FACTOR (ARF) 7 and ARF19 module (Fukaki *et al*., 2005; Okushima *et al*., 2007). The *rlf* loss of function mutants of *RLF*, show a marked reduction in the number of LRs while their ARF7/19-mediated auxin signaling is not affected, suggesting that RLF regulates LR formation independently of auxin response (Ikeyama *et al*., 2010). In addition, the *rlf* mutations cause aberrant organ development, including reduced primary root growth and decreased leaf size (Ikeyama *et al*., 2010). While the study of *RLF* in *A. thaliana* shed light on the significance of Cytb5-HBD proteins for plant growth and development, a crucial gap remains in our understanding of how Cytb5-HBD proteins orchestrate growth and development of plant organs. First, it is not known whether RLF binds to heme and works as HBP *in planta*. Second, there is limited understanding of whether Cytb5-HBD proteins share a common role in regulating organ development in land plants. To address these issues, we extend our investigation of RLF to the model bryophyte species, *Marchantia polymorpha* (Ishizaki et al., 2016; Fernandez-Pozo et al., 2022) in addition to *A. thaliana*.

In this study, we demonstrate that the heme-binding ability of RLF is necessary for its biological function. In addition, we unveil the role of Mp*RLF*, an ortholog of *RLF* in *M. polymorpha*, in orchestrating proper vegetative and reproductive development. Our findings highlight the crucial role of the Mp*RLF* gene in plant organ development. Notably, the similarities between MpRLF and the *Arabidopsis* RLF suggest the existence of a conserved mechanism, governed by RLF that intricately regulates organ development across diverse plant lineages.

## Materials and Methods

### Plant materials and growth conditions

*Marchantia polymorpha* accessions Takaragaike-1 (Tak-1; male) and Takaragaike-2 (Tak-2; female) were used as the wild-type in this study (Ishizaki *et al*., 2008). *M. polymorpha* plants were generally cultured on 1/2-strength Gamborg’s B5 medium containing 1% (w/v) agar under continuous light (50 ∼ 60 µmol m^-2^ s^-1^) at 20℃. The thalli were imaged using a dissecting microscope and measured using the Fiji imaging software (Schindelin *et al*., 2012). Observations of gemma cups and reproductive organs were performed on cultured thalli of *M. polymorpha* using a digital microscope (VHX-5000, KEYENCE, Japan). For scanning electron microscopy, thalli were frozen in liquid nitrogen and observed with a scanning electron microscope (VHX-D500, KEYENCE, Japan).

Columbia (Col) was used as the wild-type accession of *Arabidopsis thaliana* (L.) Heynh. The *rlf* mutant and *RLFpro:RLF-GFP* transgenic line have been described previously (Ikeyama *et al*., 2010). Seeds were germinated under sterile conditions on Murashige & Skoog (MS) medium containing 1% (w/v) sucrose solidified with 0.5% (w/v) gellan gum, as described previously (Goh *et al*., 2012). *A. thaliana* plants were grown at 23°C under continuous light (40 ∼ 50 µmol m^-2^ s^-1^). The number of lateral roots (LRs) and root length were determined using a dissecting microscope and the Fiji imaging software (Schindelin *et al*., 2012).

All data are the mean values for each plant considered. Experiments were performed three times, and similar values were obtained in each experiment. Statistical significance of the data was calculated using the Tukey-Kramer test.

### Constructions of vectors and generation of transgenic *A. thaliana*

Primers used for plasmid construction are listed in Supplemental Table **S2**. To generate *RLFpro:RLF-GFP, RLFpro:RLF(H161G)-GFP*, *RLFpro:RLF(H184G)-GFP*, and *RLFpro:RLF(H161G/H184G)-GFP* constructs, the promoter region of *RLF* (*RLFpro*) (804 bp) was amplified from *A. thaliana* wild-type genomic DNA using IF-pGWB501RLFpro-F and IF-pGWB501RLFpro-R primers, and cloned into pGWB501 using In-Fusion technology (Takara Bio Inc., Japan; pGWB501 *RLFpro*). Using previously synthesized coding sequence of full-length *RLF* (*RLFcds*) (Ikeyama *et al*., 2010) as a template, the *CACC-RLFcds-XhoI* fragment was amplified using CACC-RLF-F and RLF-XhoI-R primers, and cloned into the pENTR/D-TOPO cloning vector (Life Technologies, Rockville, MD, USA) to produce the pENTR-*RLF-XhoI* vector. The XhoI-GFP-SalI fragment was obtained from pBluescript II SK(+) GFP by restriction enzyme treatment and ligated with restriction enzyme-cleaved pENTR-*RLF-XhoI* to generate pENTR-*RLF-GFP*. pENTR*-RLF(H161)-GFP* and pENTR-*RLF(H184)-GFP* fragments were created by PrimeSTAR Max (Takara Bio Inc., Japan) using H161G-F/H161G-R and H184G-F/H184G-R primers, respectively. In addition, the pENTR-*RLF(H161G/H184G)-GFP* fragment was created in the same way using H184G-F/H184G-R primers with pENTR-*RLF(H161)-GFP* as the template. These entry vectors and pGWB501-*RLFpro* were treated with the Gateway LR clonase II enzyme mix (Thermo Fisher Scientific, USA) to generate *RLFpro:RLF-GFP*, *RLFpro:RLF(H161G)-GFP*, *RLFpro:RLF(H184G)-GFP*, and *RLFpro:RLF(H161G/H184G)-GFP* constructs, respectively. To generate the *RLFpro:*Mp*RLF*, pGWB501-*RLFpro* and pENTR-Mp*RLFcds* vectors (see below; Construction of vectors and generation of transgenic *M. polymorpha*) were reacted with Gateway LR clonase II enzyme mix (Thermo Fisher Scientific, USA). These binary vectors were transformed into the *rlf-1* mutant using the floral dip method (Clough & Bent, 1998) with *Agrobacterium tumefaciens* (strain C58MP90).

### Constructions of vectors and generation of transgenic *M. polymorpha*

Primers used for plasmid construction are listed in Supplemental Table S2. To generate the *Mp4g04390/*Mp*RLFpro:GUS* construct, the promoter region of Mp*RLF* (Mp*RLFpro*) was amplified by PCR from Tak-1 genomic DNA as a 4,411 bp sequence containing the start codon in the first exon using the primers MpRLFpro-CACC-F and MpRLFpro-R. The amplified fragment was cloned into the pENTR/D-TOPO cloning vector (Life Technologies, Rockville, USA) to generate the pENTR-Mp*RLFpro* vector. The Mp*RLFpro* region of the pENTR-Mp*RLFpro* vector was introduced into the Gateway binary vector pMpGWB104 (Ishizaki *et al*., 2015) using the Gateway LR clonase II Enzyme mix (Thermo Fisher Scientific, USA) to generate the Mp*RLFpro:GUS* binary construct. The Mp*RLFpro:GUS* vector was introduced into regenerating thalli of Tak-1 using *A. tumefaciens* GV2260 as previously described (Kubota *et al*., 2013). Transformants were selected using 100 µg/ml hygromycin B and 100 µg/ml cefotaxime.

Loss-of-function mutants of Mp*RLF* (Mp*rlf^ge^*) were generated using the CRISPR/Cas9 system as described previously (Sugano & Nishihama, 2018; Sugano *et al*., 2018).We selected the target sequence at the junction of the second intron and the third exon of Mp*RLF*. Synthetic oligonucleotides for the target sites shown in Table S2 were annealed, inserted into BsaI-digested pMpGE_En03 using T4 DNA Ligase (Takara Bio Inc., Japan), and then introduced into the destination vector pMpGE010 or pMpGE011 using the Gateway LR clonase II Enzyme mix (Sugano & Nishihama, 2018; Sugano *et al*., 2018). The vectors were introduced into regenerating thalli of Tak-1 or gemmae of Tak-2 using *Agrobacterium tumefaciens* GV2260 (Kubota *et al*., 2013; Tsuboyama *et al*., 2018), and transformants were selected using 100 µg/mL hygromycin or 0.5 µM chlorsulfuron.

For complementation of Mp*rlf^ge^*, the coding sequence (cds) of the full-length Mp*RLF* was amplified by RT-PCR using KOD-Plus-Neo (TOYOBO, Japan) with the primers MpRLFcds-CACC-F and MpRLFcds (+stop)-R. The Mp*RLF* coding sequence fragment was cloned into the pENTR/D-TOPO cloning vector (Thermo Fisher Scientific, USA) to produce pENTR-Mp*RLFcds*. To prevent CRISPR-Cas9 targeting of Mp*RLFcds,* a synonymous sequence substitution was introduced between nucleotide positions 472 and 492 (amino acids from G158 to R164) in Mp*RLFcds* as follows. pENTR-Mp*RLFcds* was used as a template, and the entire plasmid was amplified using PrimeSTAR Max (Takara Bio Inc., Japan) and the primers IF-MpRLF(com)-F and IF-MpRLF(com)-R to produce pENTR-Mp*RLFcds_resistant*, a CRISPR-Cas9-resistant Mp*RLFcds*. The Mp*RLF* promoter region (Mp*RLFpro*), including 4,408 bp upstream of the initiation codon, was amplified from Tak-1 genomic DNA by PCR using PrimeSTAR Max (Takara Bio Inc., Japan) with the primers IF-MpRLFpro-F and IF-MpRLFpro-R. The PCR-amplified product was cloned into the HindIII sites of pMpGWB301 (Ishizaki *et al*., 2015) using the In-Fusion HD cloning kit (Clontech, Mountain View, CA). The CRISPR-Cas9-resistant MpRLF coding sequence in pENTR-Mp*RLFcds_resistant* was introduced into the binary vector using the Gateway LR clonase II Enzyme mix (Thermo Fisher Scientific, USA) to generate the Mp*RLFpro:*Mp*RLFcds* (com) construct. The Mp*RLFpro:*Mp*RLFcds* (com) vector was transformed into the gemmae of the Mp*rlf^ge^* mutants using *A. tumefaciens* GV2260. Detailed methods are described in a previous study (Tsuboyama *et al*., 2018).

To generate Mp*E2Fpro:XVE>>*Mp*RLFcds*, the Mp*RLF* coding sequence was amplified by PCR using PrimeSTAR Max (Takara Bio Inc., Japan) with the primer set MpRLFcds-CACC-F and MpRLFcds(-stop)-R and cloned into the pENTR/D-TOPO cloning vector. LR reactions were performed on the C-terminal side of the GFP fusion destination vectors based on the Gateway binary vector pMpGWB168, (Ishida *et al*., 2022) and entry vectors. The Mp*E2Fpro:XVE>>*Mp*RLFcds-GFP* vector was introduced into regenerating thalli of Tak-1 as previously described (Kubota *et al*., 2013). Transformants were selected with 100 µg/ml hygromycin B and 100 µg/ml cefotaxime. To evaluate the effects of conditional overexpression of Mp*RLF*, gemmae of *MpEFpro:XVE>>MpRLF-GFP* plants was cultured for seven days on 1/2-strength Gamborg’s B5 medium plates, followed by seven days on 1/2- strength Gamborg’s B5 medium plates containing 10 µM ᵝ-estradiol (EST; FUJIFILM Wako Pure Chemical Corporation, Japan).

### GUS staining

For histological GUS activity assays, the thalli of Mp*RLFpro:GUS*/Tak-1 were de-aerated twice for 5 min, incubated in GUS staining solution (0.5 mM potassium ferrocyanide, 0.5 mM potassium ferricyanide, and 1 mM X-Gluc) at 37°C for 4 hours and later cleared with 70% (v/v) ethanol. GUS-stained samples were mounted in 6% (w/v) Agar. Sections of 10∼15 µm samples were cut using a microtome (HM325; MICROM International Inc., Walldorf, Germany). The sections were imaged by differential interference contrast microscopy (Leica DM6000 microscope, Leica Microsystems, Germany) and assembled by e-Tilling (Mitani corp., Japan).

### Confocal laser scanning microscopy

The gemmae expressing EST-inducible MpRLF-GFP were counterstained with 10 µg/ml of propidium iodide (PI) for 5 minutes and observed by confocal laser scanning microscopy using an Olympus FV1000 confocal microscope (Olympus/Evident, Japan). The 473 nm line and the 559 nm line of an LD laser were used to excite GFP and PI, respectively. Images were acquired using the UPlanSAPO 40x/0.95 objective (Olympus/Evident, Japan). Images were processed using the Fiji/ImageJ program (Schindelin *et al*., 2012).

### Multiple sequence alignments and phylogenetic analysis

The amino acid sequence of *Arabidopsis thaliana* RLF (At5g09680) was used as the template for identifying similar sequences of other plant RLF orthologs using PSI-BLAST. Ortholog sequences from eighteen different green plants were meticulously chosen based on an E-value threshold of 1e-20, and multiple sequence alignment were conducted using ClustalW (Waterhouse *et al*., 2009; Katoh & Standley, 2013). The MAFFT Multiple Sequence Alignment output was color-coded in blue to highlight conserved regions. In the construction of the molecular phylogenetic tree, amino acid sequences of RLF and 19 other plant RLFs were obtained from the MAFFT Multiple Sequence Alignment. Subsequently, a phylogenetic tree was constructed using MEGA11 to calculate the similarity between RLF and other plant RLFs: *Arabidopsis thaliana* (RLF/At5g09680), *Tarenaya hassleriana* (XP 010547268), *Cucurbita moschata* (XP 022960400), *Populus trichocarpa* (POPTR 001G263800), *Glycine max* (GLYMA 09G227600, GLYMA 12G009100), *Citrus sinensis* (KAH9774569), *Oryza sativa subsp. Japonica* (Os07g0232200), *Zea mays* (LOC100272407), *Amborella trichopoda* (XP 011624637.1), *Cryptomeria japonica* (XP 057826847.1), *Marchantia polymorpha* (MpRLF/Mp3g04390), *Marchantia paleacea* (KAG6551382.1), *Physcomitrium patens* (Pp3c2023480), *Ceratopteris richardii* (KP509 01G108300), *Selaginella moellendorffii* (SELMODRAFT 127066, SELMODRAFT 138045), *Chara braunii* (CBR g29320), *Chlamydomonas reinhardtii* (CHLRE 13g574800v5), *Volvox reticuliferus* (Vretifemale 5060). In this process, the bootstrap method was performed 1000 times and the WAG model (Whelan & Goldman, 2001) was applied.

### Protein purification and measurement of absorbance spectra

Purification of recombinant proteins was performed according to the protocol of the pMAL Protein Fusion and Purification System (New England Biolabs, Japan). The expression clone Maltose Binding Protein (MBP)-RLF was obtained by an LR reaction of the pMal-c5X Vector, in which contains the *att*R1 sequence, *ccdB* gene, and *att*R2 sequence at the multicloning site, with RLFcds/pENTR. To generate MBP-mutant RLF, we constructed *RLFcds(H161A)*/pENTR, *RLFcd(H184A)*/pENTR by using *RLFcds*/pENTR as template with H161A-F/H161A-R and H184A-F/H184-R primers using PrimeSTAR Max (Takara Bio Inc., Japan). To construct *RLFcd(H161A/H184A)/*pENTR, *RLFcds(H161A)*/pENTR was used as a template using H184A/-F/H184-R primers with PrimeSTAR Max (Takara Bio Inc., Japan). These mutant *RLF/*pENTR were used in the same way to perform LR reactions to construct *MBP-RLF(H161A), MBP-RLF(H184A)* and *MBP-RLF(H161A.H184A)*. The expression clone was transformed into *E. coli* JM109. The growing colonies were pre-cultured in 5 ml of LB liquid medium containing 50 µg/ml ampicillin and 0.2% (w/v) glucose at 37°C for 18 h. Three ml of the pre-culture medium was added to 300 ml of LB liquid medium containing 50 µg/ml ampicillin and 0.2% (w/v) glucose at 37°C, 120 rpm, 2 ∼3 hours of incubation, followed by shaking at 20°C, 120 rpm for 1 hour. 1 ml of 100 mM Isopropyl β-D-thioglactopyranoside (IPTG, Takara Bio Inc., Japan) was added to the culture medium, and the mixture was shaken at 20°C, 90 rpm for 20 h. The cell pellet was collected by centrifugation at 4°C, 10 min, and 3,600 rpm, resuspended in 1 x PBS and centrifugation at 4°C, 10 min, and 3,600 rpm. The collected cell pellet was stored at -80°C.

Upon use, the pellet was suspended in 5 ml of Buffer A (20 mM Tris-HCl pH 7.5, 200 mM NaCl) and transferred to a 10 ml beaker. To suppress the activity of proteolytic enzymes, 20 µl of 10 mg/ml PMSF was added to the bacterial suspension prior to lysis by sonication. After treatment, the extract was aliquoted into 1.5 ml tubes, centrifuged at 4°C for 15 minutes at 16,000 × g, and the supernatant was collected in 50 ml tubes. 300 µl of Amylose Resin (50% (w/v) slurry) was added and allowed to stand at 4°C for 30 minutes with moderate mixing. The mixture was then centrifuged at 4°C for 1 minute at 500 × g, and the supernatant was discarded. Subsequently, 1 ml of Buffer A was added to the resin precipitate and collected in a 2 ml tube. After washing, 250 µl of Buffer B (20 mM Tris-HCl pH 9.5, 200 mM NaCl, 10 mM maltose) was added to the resin with the supernatant discarded and allowed to stand at 4°C for 30 minutes with moderate mixing. After centrifugation at 4°C, 1 min, 500 × g, the supernatant was collected, and 6 x Stock buffer (60% (v/v) glycerol, 0.3% (v/v) Triton-100) was added prior to storing at -80°C.

For measuring the ultraviolet (UV)-visible absorption spectrum of RLF proteins, samples were centrifuged to remove impurities and exchanged by ultrafiltration into a buffer containing 20 mM Tris-HCl (pH 8.0), 500 mM NaCl, and 6% (v/v) glycerol. The protein concentration was adjusted to 1.5 µM. Excess sodium ferricyanide in solution and a few grains of solid dithionite were added to samples, to establish the oxidized and reduced heme conditions respectively. Measurements of the absorbance spectra were performed using a spectrophotometer UV-1900i (SHIMADZU, Japan).

### RNA-seq and transcriptomic analysis

Total RNA was purified from 16-day-old thalli (approximately 100 mg fresh weight) using the RNeasy Plant Mini Kit (Qiagen); RNA was quantified using a NanoDrop ND1000 spectrophotometer (Thermo Fisher Scientific, USA), RNA aliquots of 10 µg/ml or greater from three biological replicates were used for RNA-Seq analysis. Library preparation and RNA-seq analysis were contracted to BGI JAPAN (URL https://www.bgi.com/jp/home), and clean reads were obtained from each library. The resulting reads were mapped using MpTak_v6.1r2 from Marpolbase (URL https://marchantia.info/), excluding sequences on the U chromosome from the data. We used HISAT2 (Kim *et al*., 2015) to align the clean reads to the reference genome, and then StringTie (Pertea *et al*., 2015) to assemble them. Finally, the assembly results of all samples were combined using Cuffmerge. We used Bowtie2 (Langmead & Salzberg, 2012) to align clean reads to the reference sequence, and then use RSEM (Li & Dewey, 2011) to calculate the expression of genes and transcripts. Enrichment analysis uses Gene Ontology (GO) and KEGG to classify gene function and evaluate gene abundance within a particular functional module. Statistically, a hypergeminia model calculates the p-value for each functional module. False discovery rate (FDR) correction is performed to determine significant enrichment; functional modules with an FDR of 0.01 or less are considered significant. The bubble chart and GO hierarchical display are based on the results of GO enrichment analysis using GOATOOLS (Klopfenstein *et al*., 2018). Each bubble indicates the number of genes associated with the GO term, and the color represents the enrichment score (-log10 (p_fdr_bh)). -log10 (p_fdr_bh) is the value resulting from a logarithmic transformation of the value of p_fdr_bh with a base of 10 and sign inversion. The value p_fdr_bh is calculated by considering multiple comparisons using the Benjamin-Hochberg method. This method is applied to p_uncorrected, which indicates whether each GO term is significantly enriched. The GO hierarchy display visualizes the relationship between each GO term using Graphviz on the results of the GO enrichment analysis (Ellson *et al*., 2002; Klopfenstein *et al*., 2018). Each GO term includes the log-transformed value of p_uncorrected.

## Results

### Heme-binding ability is necessary for RLF function in *A. thaliana*

Two histidine residues in cytochrome *b*_5_-like heme-binding domain (Cytb5-HBD) are known to be important for heme binding (Cowley *et al*., 2002; Schenkman & Jansson, 2003). Comparison of the amino acid sequence of the Cytb5-HBD in RLF with those of the other Cytb5-HBD proteins in *A. thaliana* revealed that two histidine residues at positions 161 and 184 of RLF correspond to the predicted heme-binding sites and are conserved among all the Cytb5-HBD proteins (**Figs 1a**, **S1a**). To confirm the importance of these histidine residues, we purified recombinant wild-type and mutant RLF proteins with individual or combined histidine-to-alanine substitutions (H161A, H184A, and H161A/H184A) and measured their absorbance spectra. The oxidized form of wild-type RLF showed a peak in the soret band (413 nm) characteristic of Cytb5-HBD proteins. In contrast, the reduced form of RLF showed peaks in the αβ band (527 and 557 nm) in addition to the soret band (424 nm) (**Figs 1b**, **S1b,c**). These absorption spectra of wild-type RLF are characteristic of the Cytb5-HBD proteins (Smith *et al*., 1994; Wahl *et al*., 2010). In contrast, the spectra of the mutant RLF proteins (H161A, H184A, and H161A/H184A) did not show any peaks. The structure of each mutant RLF protein, predicted using ColabFold, showed that the C-terminal CytB5-HBD is highly conserved between wild-type and mutant proteins (**Fig. S1b**; Mirdita *et al*., 2022). This suggests that the heme-binding site substitutions do not affect the overall protein structure. These results indicate that RLF is a heme-binding protein and confirm that two histidine residues at positions 161 and 184 are necessary for heme-binding to RLF *in vitro*.

**Fig. 1.**
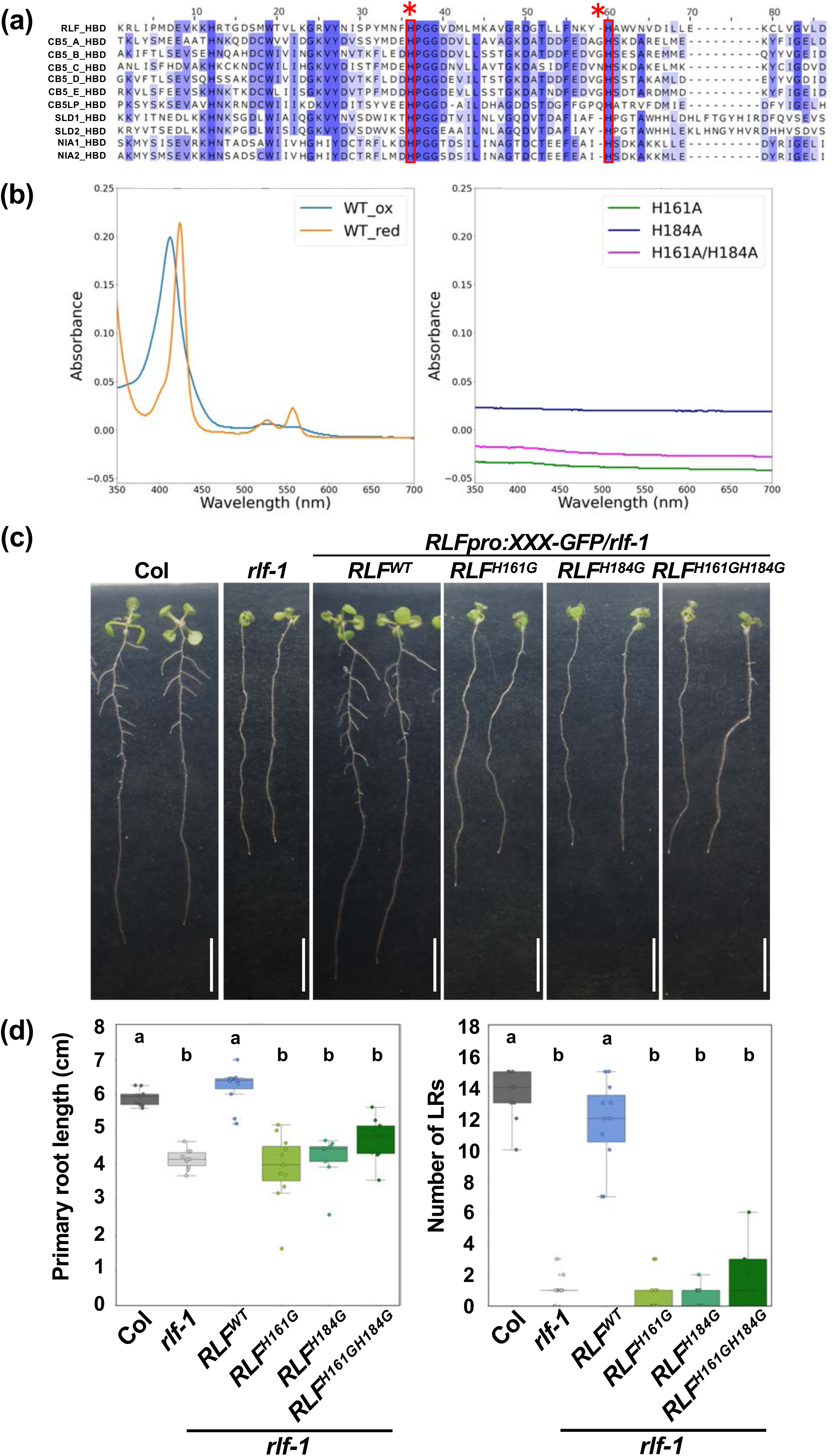
RLF functions as a heme-binding protein (**a**) Alignment of the Cytb5-HBD domain among Cytb5-HBD proteins in *Arabidopsis thaliana*. Histidine residues at positions 161 and 184 of the RLF protein, indicated by asterisks, and are predicted to be heme-binding sites. (**b**) Absorbance spectra of wild-type and mutant MBP-RLF proteins. The measurement method is described by (Takahashi *et al*., 2008). Rescue of the *rlf* phenotype (**c**) and primary root length and number of lateral roots phenotypes of *rlf* (**d**) by *RLF*, *RLF^H161G^*, *RLF^H184G^ and RLF^H161GH184G^* using RLF-GFP fusions under the control of the *RLF* promoter in 9-day-old seedlings. Scale bar = 1 cm. (9 ≦ n ≦ 11). Samples denoted by different letters are significantly different (*P* < 0.05, Tukey‒Kramer test).

Although RLF binds to heme *in vitro*, the functional role of heme binding to RLF *in planta* remains unclear. To address this question, we generated transgenic *A. thaliana* plants expressing the mutant RLF where histidine residues at positions 161 and 184, individually or in combination, were replaced with glycine (H161G, H184G, and H161G/H184G), in the *rlf-1* mutant background (Ikeyama *et al*., 2010). As previously shown, the *rlf-1* mutant displayed a marked reduction in the number of lateral roots (LRs) and reduced primary root growth, compared to the wild type (**Fig. 1c,d**). The transgenes expressing the GFP-tagged mutant RLF [*RLFpro:RLF(H161G)-GFP*, *RLFpro:RLF(H184G)-GFP*, and *RLFpro:RLF(H161G/H184G)-GFP*] under the control of the *RLF* promoter failed to restore LR formation and primary root growth in the *rlf-1* mutant. In contrast, the *RLFpro:RLF-GFP* transgene expressing the GFP-tagged wild-type RLF successfully rescued the *rlf-1* mutant phenotype (**Fig. 1c,d**). These results indicate that the two histidine residues at positions 161 and 184 are important for RLF function *in planta*, implying that heme binding is necessary for the biological function of RLF.

### Conservation of RLF across diverse plant species

A previous study has shown that the *RLF* orthologous genes are found in several plants, including monocot rice (*Oryza sativa*) and moss (*Physcomitrium patens*) (Ikeyama *et al*., 2010). To confirm amino acid sequence conservation among RLF orthologs, we first performed multiple alignment comparisons using Jalview (Waterhouse *et al*., 2009) on the amino acid sequences from *Arabidopsis thaliana* (At5g09680), *Zea mays* (LOC100272407), *Oryza sativa* (Os07g0232200), *Cryptomeria japonica* (XP_057826847), *Marchantia polymorpha* (M3g04390), *Physcomitrium patens* (Pp3c20_23480), *Ceratopteris richardii* (KP509_01G108300), *Chlamydomonas reinhardtii* (Cre13_g 574800), *Volvox reticuliferus* (Vretifemale_5060). This analysis revealed that the N-terminal Cytb5-HBD region is highly conserved among RLF orthologs (**Fig. S2**), while the length and conservation of the disordered region on the N-terminal side vary significantly among these orthologs (**Fig. S2**). Particularly, the two histidine residues, that are part of the heme-binding site, are highly conserved in the N-terminal part of the CytB5-HBD region (**Fig. S2**). These findings suggest that the heme binding is crucial for the function of RLF orthologs.

We further constructed the molecular phylogenetic tree to examine how *RLF* orthologs are conserved in the plant kingdom (**Fig. S3**). These analyses demonstrated that *RLF* orthologs are conserved across a wide variety of plants, including angiosperms, gymnosperms, ferns, liverworts, mosses, charophyte algae, and chlorophyta. Furthermore, most plant species have a single *RLF* ortholog, with the exception of *Glycine max* and *Selaginella moellendorffii* (URL https://phytozome-next.jgi.doe.gov/; **Fig. S3**).

### Mp*RLF*, the *RLF* ortholog in *M. polymorpha*, complements the *rlf* mutant phenotype in *A. thaliana*

Although *RLF* orthologs are conserved among plants, we do not know whether the mechanisms of RLF-dependent organ development are also conserved across species. To examine whether these *RLF* orthologs also regulate organ development, we used the model bryophyte species *M. polymorpha*, and named the *M. polymorpha RLF* ortholog as Mp*RLF.* First, to assess whether Mp*RLF* gene function is preserved between *M. polymorpha* and *A. thaliana*, we expressed Mp*RLFcds* under the control of the *RLF* promoter in the *A. thaliana rlf* mutant background (*RLFpro:*Mp*RLFcds/rlf-1*). As shown in **Figure 2**, expression of MpRLF in *rlf* mutant fully restored the reduction of LR number and primary root length phenotypes (**Fig. 2**). These results indicate that MpRLF is functionally interchangeable with *Arabidopsis* RLF, suggesting that the mechanisms of RLF-dependent organ development are conserved between *A. thaliana* and *M. polymorpha*.

**Fig. 2.**
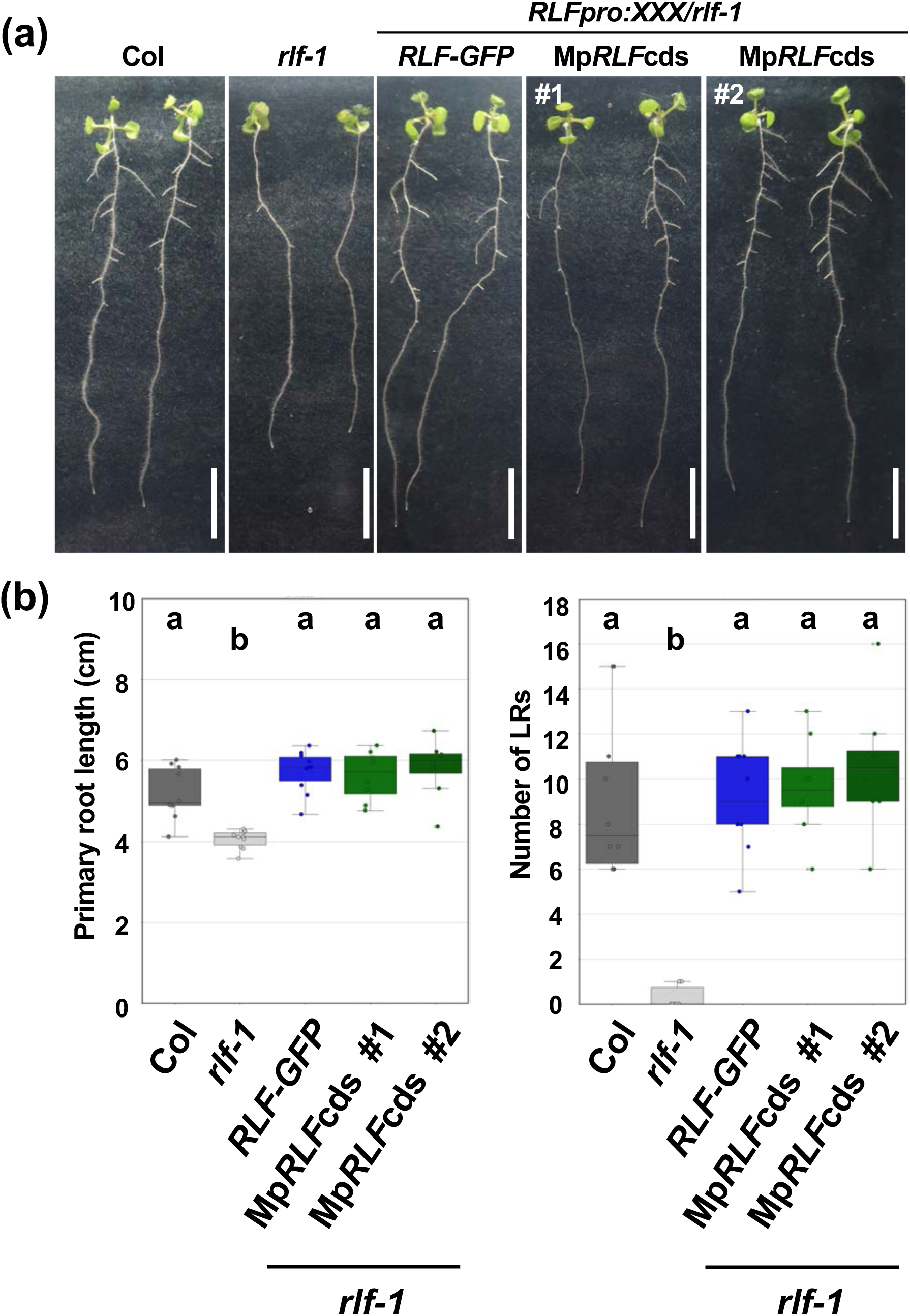
Mp*RLF* complements the *rlf* mutant phenotype in *A. thaliana* (**a**) 9*-*day-old seedlings of wild type, *rlf-1* mutant, and *rlf-1* mutants expressing *RLFpro:RLFcds-GFP* and *RLFpro:*Mp*RLFcds*. Scale bar = 1 cm. (**b, c**) Primary root length and number of LRs in 9 day-old wild-type (Col), *rlf-1, RLFpro::RLF-GFP/rlf-1* and *RLFpro::*Mp*RLFcds/rlf-1* seedlings (8 ≦ n ≦ 10). Samples denoted by different letters are significantly different (*P* < 0.05, Tukey‒Kramer test).

### Mp*RLF* is expressed along the midrib of the thallus in *M. polymorpha*

Next, we examined the expression pattern of Mp*RLF* in *M. polymorpha.* The recently published *M. polymorpha* database, Marpolbase Expression (URL https://marchantia.info/mbex/; Kawamura et al., 2022), suggests that Mp*RLF* is expressed in various organs such as mature antheridiophores, antheridia, archegoniophore, archegonia and gemma cups (**Fig. S4**). To elucidate the expression pattern of Mp*RLF* in detail, we generated transgenic *M. polymorpha* plants expressing the GUS reporter gene under the control of the Mp*RLF* promoter (Mp*RLFpro:GUS*). In Mp*RLFpro:GUS* lines, GUS activity was detected, by histological staining, in the midrib and near the apical meristem of gemmae and 14-day-old thalli (**Fig. 3a**). In transverse sections of the mature thallus, GUS activity was detected along the midrib on the ventral side of the thallus. In longitudinal sections, GUS activity was detected at the bottom of the gemma cup **(Fig. 3b,c**). In contrast, GUS activity was absent in the rhizoid. These results indicate that Mp*RLF* is expressed in the thallus and during gemma formation.

**Fig. 3.**
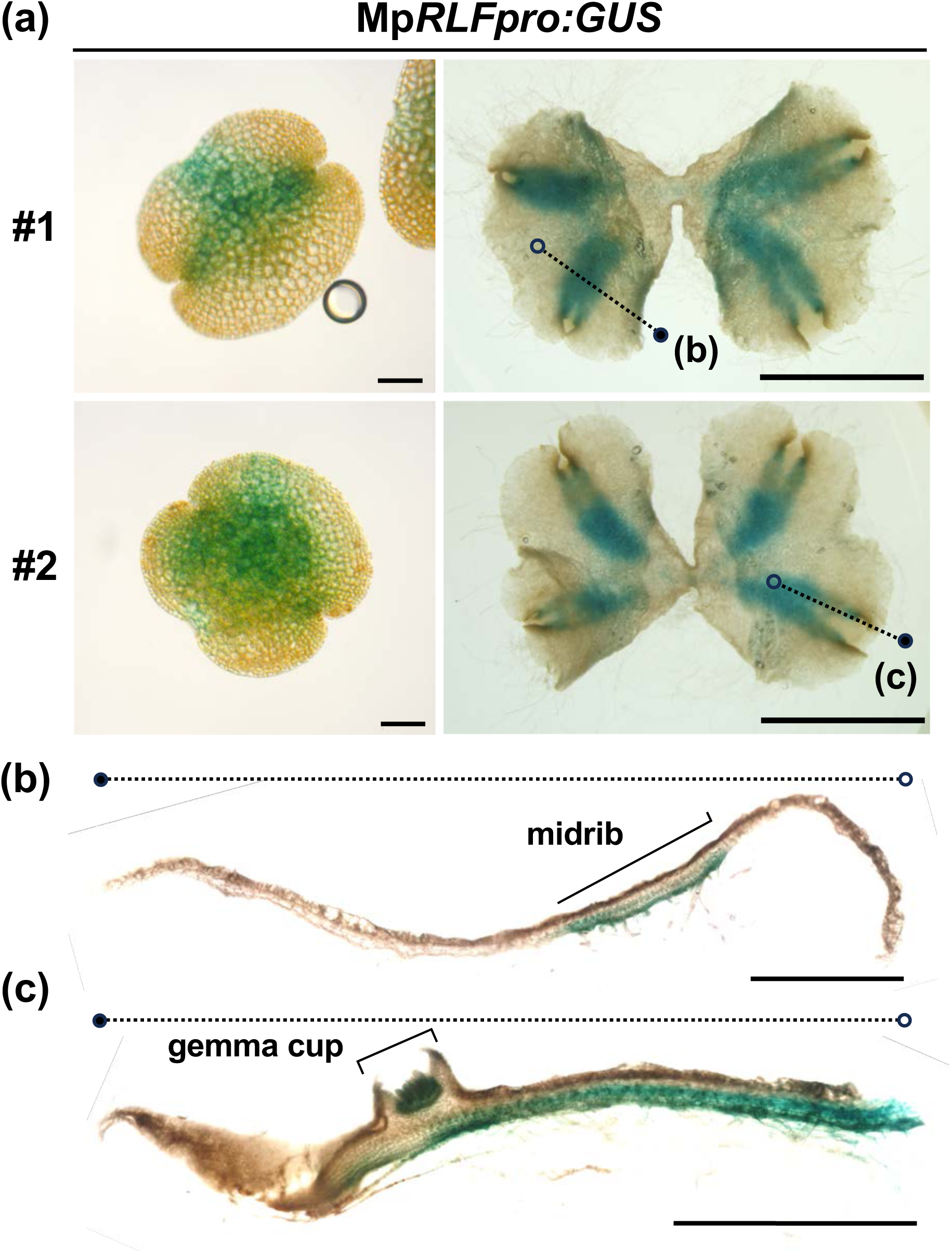
Expression pattern of Mp*RLF* in *M. polymorpha* (**a**) GUS staining of Mp*RLFpro:GUS* reporter lines. Left panel shows gemma stained for 24 hours and right panel shows thallus stained for 4 hours after 14 days of growth (left panels: scale bar = 100 µm, right panels: scale bar = 5 mm). Dashed lines indicate directions of cross sections shown in (**b**) and (**c**). (**b, c**) Expression pattern of Mp*RLFpro:GUS* in cross sections of thallus. These images show a transverse section (**b**) and a longitudinal section (**c**) of thallus stained with GUS for 4 h after 14 days of growth. Scale bar = 1 mm.

### Mp*RLF* is essential for proper thallus and gemma cup formation

To investigate the biological role of Mp*RLF*, we produced a loss-of-function mutant of Mp*RLF* (Mp*rlf^ge^*) using the CRISPR-Cas9 method. The target sequence of CRISPR-Cas9 was designed at the junction of the second intron and the third exon, upstream of the Cytb5-HBD (**Fig. 4a**). All three alleles of the Mp*rlf^ge^* mutant obtained exhibited a frameshift leading to a premature stop codon and showed growth retardation compared to wild type (**Figs 4b-d**, **S5a**). The wild-type plant formed the gemma cup with gemmae on the dorsal surface of the thallus, while in all three Mp*rlf^ge^* mutants, the formation of proper gemma cups was inhibited and gemma-like structures formed directly on the thallus surface (**Fig. 5a**). Under far-red light enriched illumination, the wild type forms an antheridiophore containing antheridia within, whereas the mutants formed exposed antheridium-like structures on the surface of the antheridiophore (**Fig. 5b**). In addition, when the Mp*rlf^ge^*mutant was generated in a female Tak-2 background, the archegoniophore appeared to be smaller than in the wild type (**Figs 5b, S5c**). A complementation test with an Mp*RLF* coding sequence with synonymous CRISPR-resistant mutations (Mp*RLFcds_resistant*) showed that transgenic plants expressing Mp*RLFcds_resistant* under the control of the Mp*RLF* promoter in an Mp*rlf^ge^* background grew as well as the wild type (**Fig. S5b**). These results indicate that phenotypes observed in Mp*rlf^ge^* (retardation of thallus growth and impaired gemma cup formation) are due to the mutation in the Mp*RLF* gene, thereby confirming that Mp*RLF* is essential for proper thallus growth and gemma cup formation.

**Fig. 4.**
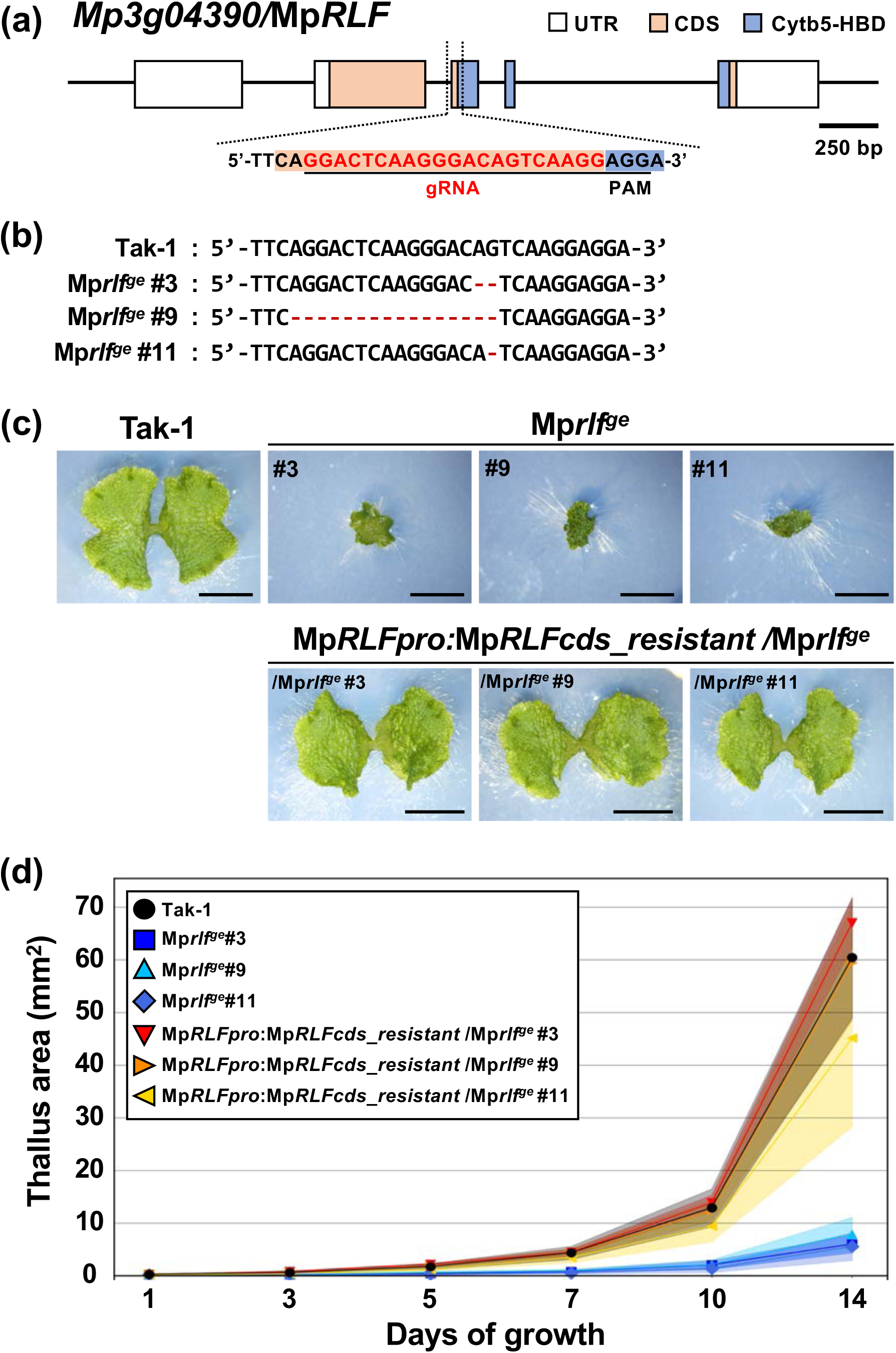
Mutations in Mp*RLF* inhibit vegetative growth and development in *M. polymorpha* (**a**) *Mp3g04390/*Mp*RLF* gene structure and selected target sequence for CRISPR mutagenesis. White, orange, and blue boxes indicate exons in the untranslated region, protein coding region, and Cytb5-HBD, respectively. The underlined sequence is the CRISPR’s target sequences; red bases indicate gRNA sequence and black bases indicate PAM (Protospacer Adjacent Motif) sequences. (**b**) Sanger sequencing analysis of mutations at gRNA target site. Dashed lines indicate deleted bases. (**c**) Wild type *M. polymorpha* (Tak-1), Mp*rlf^ge^* mutants, and complementation lines grown for 10 days. Scale bar = 5 mm. (**d**) Thallus growth area for 2 weeks after gemma germination. Data are presented as means, with confidence intervals filled in for each data (95%) (8 ≦ n ≦ 10).

**Fig. 5.**
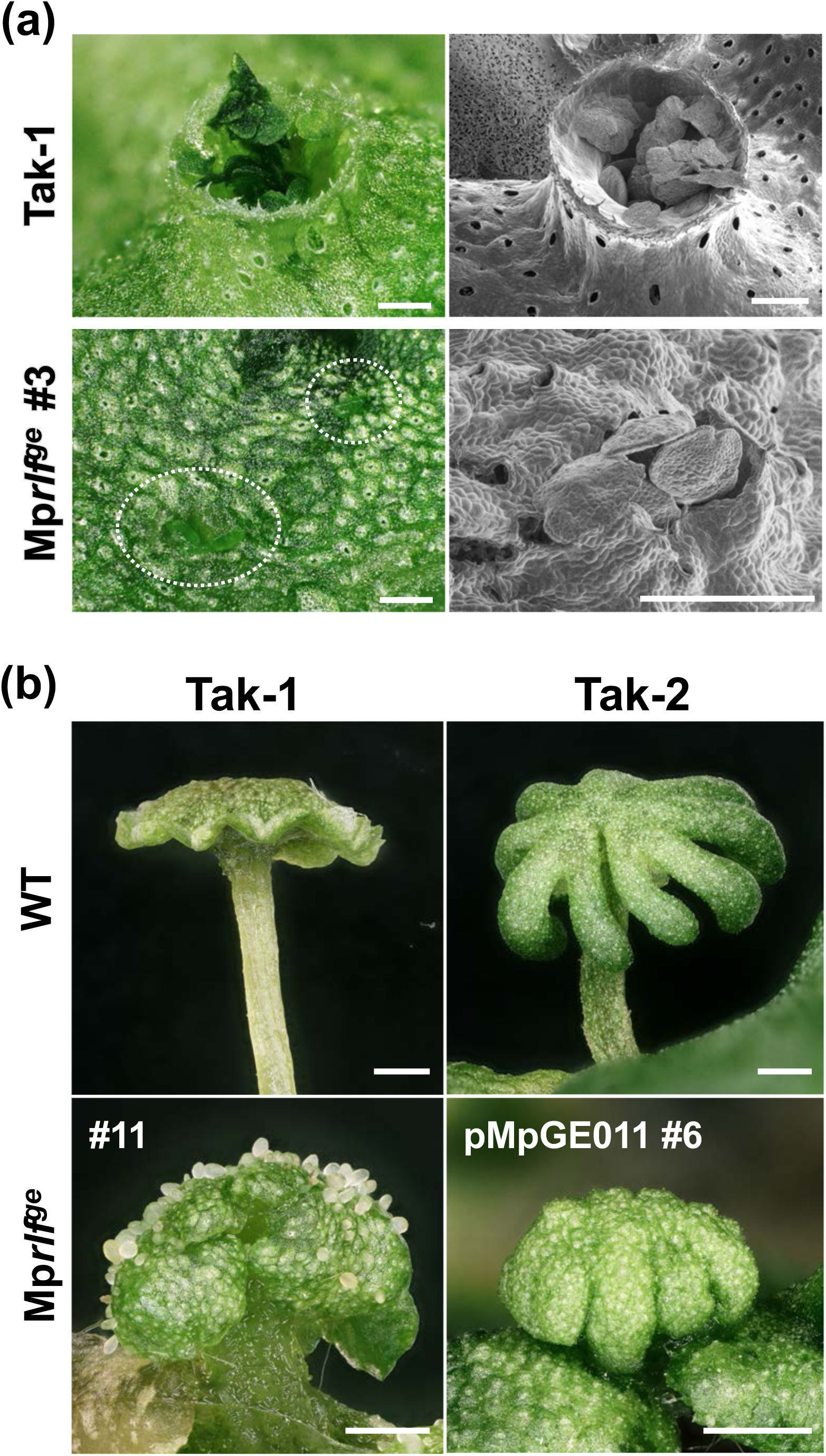
Mutations in Mp*RLF* inhibit the formation of gemma cups and reproductive organs (**a**) Dorsal surface of the thallus of wild type *M. polymorpha* (Tak-1) (top) and Mp*rlf^ge^#3* mutant (bottom) grown for three weeks. The left panels show brightfield images; the right panels show scanning electron microscope images. White dotted circles indicate gemmae without gemma cups. Scale bars = 1 mm. (**b**) Reproductive organs of wild type *M. polymorpha* and Mp*rlf^ge^* mutant. Left and right sides show male (Tak-1) and female (Tak-2) plants, respectively. The gemmae were incubated under white light for 2 weeks and then under far-red light for another 6 weeks to induce reproductive growth. pMpGE011 line #6 was used for Mp*rlf^ge^*/Tak-2 (Fig. **S5c**). Scale bar = 1 mm.

To investigate the impact of Mp*RLF* overexpression on organ development, we generated transgenic plants with Mp*RLFcds-GFP* expression inducible by estradiol (EST) under the control of *E2Fpro* (*E2Fpro:XVE>>*Mp*RLFcds-GFP*). We found that EST-induced overexpression of MpRLF (MpRLF-GFP) clearly suppressed thallus growth (**Fig. S6**), suggesting that proper regulation of Mp*RLF* expression is important for vegetative growth of *M. polymorpha*. Taken together, these results indicate that Mp*RLF* is required for proper organ formation during vegetative and reproductive growth of *M. polymorpha*.

### MpRLF protein is localized to the cytosol

In *A. thaliana*, RLF localizes to the cytosol (Ikeyama *et al*., 2010). To investigate the subcellular localization of MpRLF, the *E2Fpro:XVE>>*Mp*RLFcds-GFP* lines described above were used for observation. In the epidermal cells near the apical notch of the *E2Fpro::XVE>>*Mp*RLFcds-GFP* gemmaling grown for 24 h on EST containing medium, MpRLF-GFP was localized to the cytosol and did not co-localize with chloroplasts, the nucleus, or the plasma membrane (**Fig. 6**), which is consistent with the subcellular localization of the *Arabidopsis* RLF (RLF-GFP) protein (Ikeyama *et al*., 2010). These results suggest that MpRLF functions in the cytosol.

**Fig. 6.**
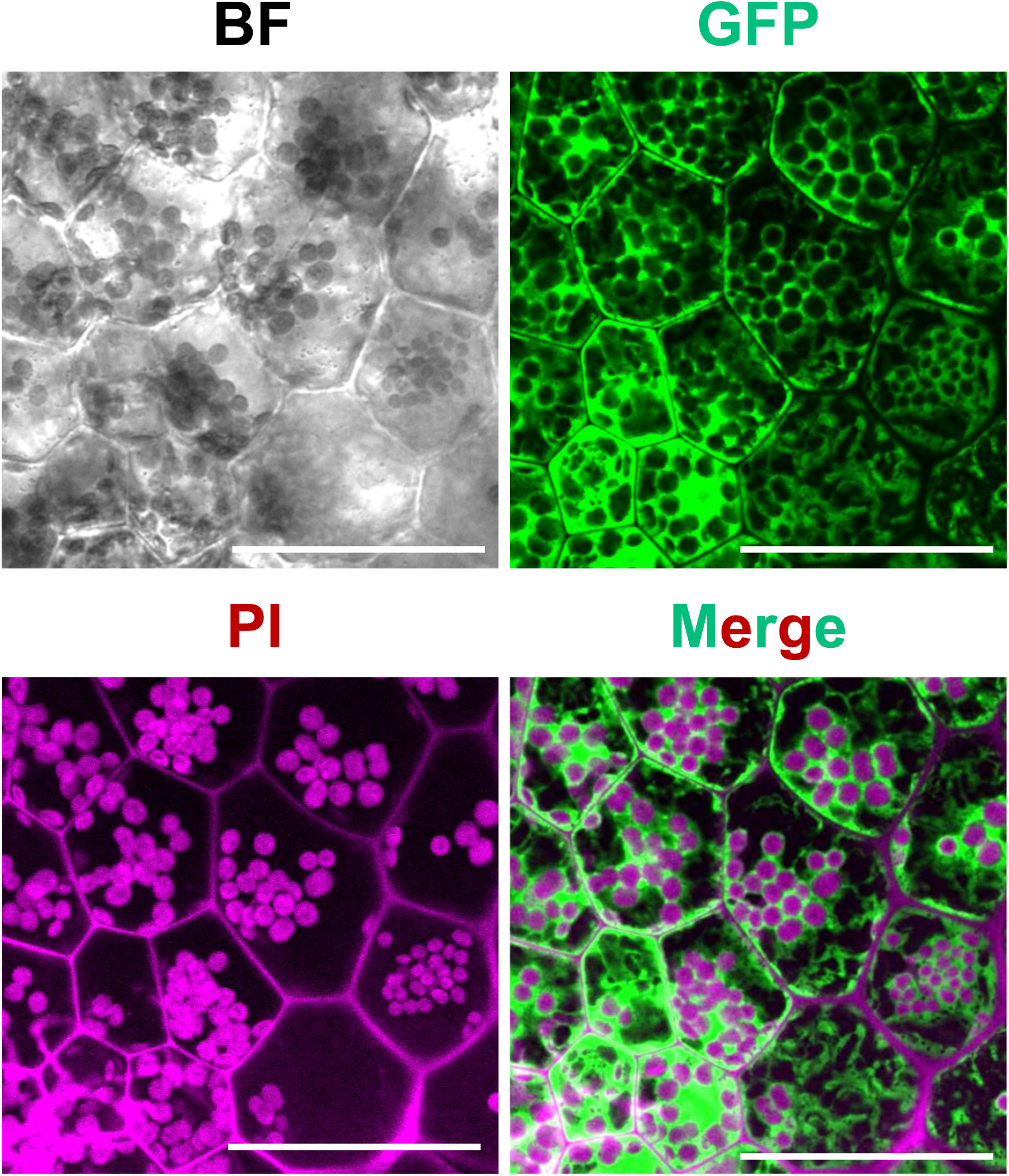
MpRLF-GFP is localized to the cytosol Gemmae of *E2Fpro::XVE>>*Mp*RLFcdc-GFP/*Tak-1 were treated with 10 µM estradiol for 24 h. Green indicates GFP fluorescence of MpRLFcds-GFP, and magenta indicates chloroplast autofluorescence and PI fluorescence. Scale bar = 50 µm.

### MpRLF strongly alters expression of metabolism-related genes

Finally, to gain insight into the pathways through which MpRLF regulates organ growth and development, we performed RNA sequencing (RNA-Seq) analysis using Tak-1 and Mp*rlf^ge^* mutant (Mp*rlf^ge^*#3). Of the sequences obtained from the clean reads, 97.6% could be mapped to the *M. polymorpha* and *M. polymorpha subsp. ruderalis* genome (Marpolbase MpTak_v6.1r2; (Bowman *et al*., 2017; Montgomery *et al*., 2020)) (Fig. **S7a**). There are 3,734 differentially expressed genes (DEGs) in Mp*rlf^ge^*, with 1,839 upregulated and 1895 downregulated (**Fig. 7a**).

**Fig. 7.**
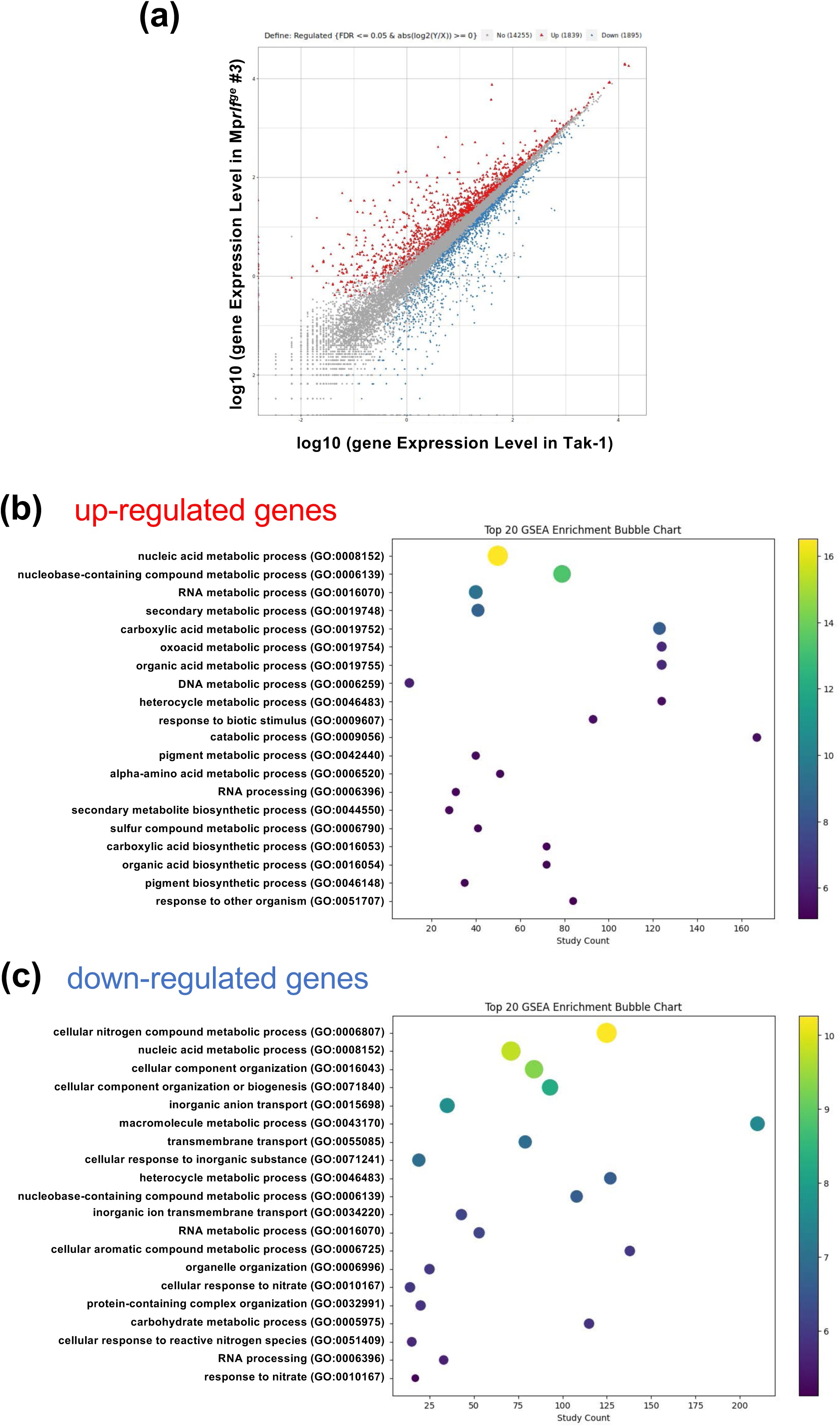
Transcriptome analysis with Mp*RLF* loss-of-function mutants (**a**) MA plot of wild type and Mp*rlf^ge^* #3 mutant. Each dot represents the log10 fold change of a gene, with red and blue dots representing up- and down-regulated genes in Mp*rlf^ge^* #3 compared to wild type, respectively. (**b, c**) Gene set enrichment analysis (GSEA) based on biological processes in the GO terms. Each bubble chart displays the top 20 GO terms, showing log10(p_fdr_bh) resulting from GO enrichment analysis for up-regulated (**b**) and down-regulated (**c**) genes in Mp*rlf^ge^* #3. The horizontal axis represents the number of genes, while the color bar represents log10(p_fdr_bh). The value - log10(p_fdr_bh) is obtained by converting the p_fdr_b value to its negative logarithm base 10.

To better characterize the gene expression variations in Mp*rlf^ge^*, we performed Gene Ontology (GO) enrichment analyses. The results indicate that the top affected biological processes in Mp*rlf^ge^* are cellular processes (GO:0009987) and metabolic processes (GO:0008152). In cellular components, the top affected processes are cell parts (GO:0044464), cell (GO:0005623), membrane (GO:0016020), membrane parts (GO 0044425), organelle (GO:0043226), and organelle part (GO:0044446). In molecular function, binding (GO:0005488) and catalytic activity (GO:0003824) are the top induced processes (**Fig. S7b**). Furthermore, KEGG (Kyoto Encyclopedia of Genes and Genomes) pathway analysis of differentially expressed genes showed they are enriched for metabolism-related terms (**Fig. S7c**). In addition, gene set enrichment analysis (GSEA) using bubble charts showed that, among biological processes, those related to metabolic processes—such as nucleic acid metabolic processes (GO:0090304), secondary metabolic process (GO:0019748), and carboxylic acid metabolic process (GO:0019752) —are enriched in up-regulated genes (**Fig. 7b**). In the same analysis for the down-regulated genes, the cellular nitrogen compound metabolic process (GO:0034641) is particularly enriched. These findings suggest that the metabolic state is drastically altered in Mp*rlf^ge^* (**Fig. 7c**).

GSEA, based on molecular function, showed that GO terms related to oxidoreduction—such as oxidoreductase activity (GO:0016491) and catechol oxidase activity (GO:0004097)—are enriched in upregulated genes, and transmembrane transporter activity (GO:0022857) and heme binding (GO:0020037) are enriched in down-regulated genes (**Fig. S8a**). GSEA, based on cellular components, revealed that up-regulated genes are enriched in apoplast (GO:0048046) and nuclear protein-containing complex (GO:0140513), while down-regulated genes are enriched in protein-containing complex (GO:0032991) and plasma membrane (GO:0005886) (**Fig. S8b**). In addition, we performed GO hierarchical analysis of biological processes on the top 100 up-regulated and the top 200 down-regulated genes. The up-regulated genes are enriched for GO terms related to chitin metabolism (**Fig. 8a**, Table **S1**). For down-regulated genes, GO terms related to response, transport, and metabolism to nitrate are enriched (**Fig. 8b**, **Table S1**). These results suggest that the aberrant organ growth and development observed in Mp*rlf^ge^* might be caused by changes in the metabolic status, potentially involving MpRLF as a Cytb5-HBD protein.

**Fig. 8.**
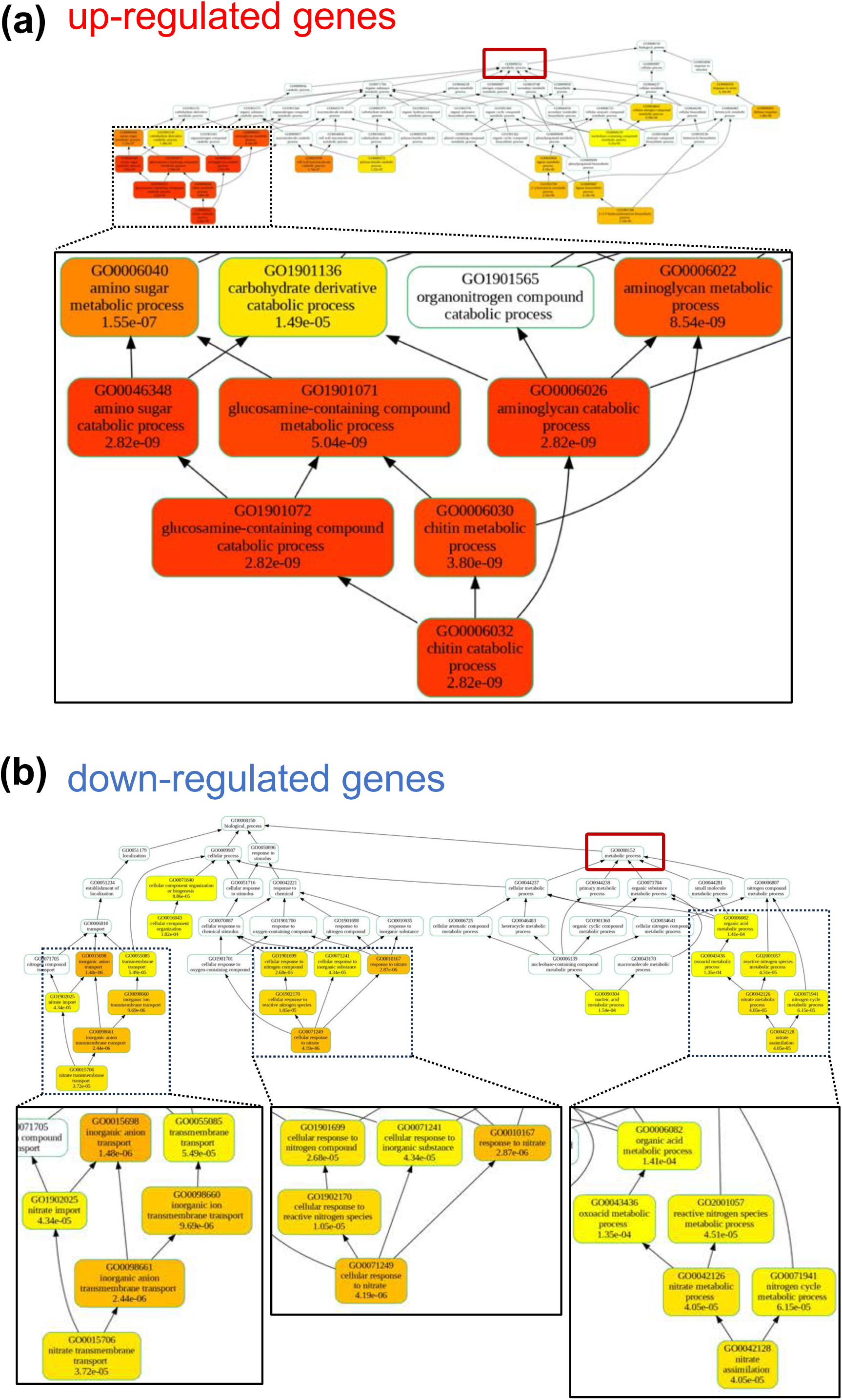
GO hierarchy of top 100 upregulated and top 200 downregulated genes GO hierarchical structure based on biological processes for the top 100 genes with upregulated expression (**a**) and the top 200 genes with downregulated expression (**b**), respectively. Red boxes indicate metabolic process (GO0008152). The GO hierarchy was created concerning by referring to Ellson et al. (2002) and Klopfenstein et al. (2018).

## Discussion

In this study, we demonstrated that *Arabidopsis* RLF is a heme binding protein and confirmed that two histidine residues at positions 161 and 184 within Cytb5-HBD are necessary for heme-binding *in vitro* and for RLF function *in planta*. *RLF* orthologs are highly conserved in various green plants and the two histidine residues, essential for heme binding within Cytb5-HBD, are found in RLF orthologs. These results imply that RLF orthologs in other plants also bind to heme via two histidine residues and that the heme-binding ability of Cytb5-HBD is necessary for the biological function of RLF proteins. Furthermore, we showed that Mp*RLF*, a *RLF* ortholog in *M. polymorpha*, rescues the *rlf* the LR formation mutant phenotype in *A. thaliana* and that Mp*RLF* is crucial for organ development in *M. polymorpha*. These results suggest that RLF orthologs in the other plants may also play a role in organ development. Therefore, it will be interesting to verify whether RLF orthologs from the different plants are able to complement either the *rlf* mutant in *A. thaliana* or the Mp*rlf^ge^* mutant in *M. polymorpha*, or both.

On the other hand, the N-terminal disordered region of RLFs varies among different plant species (**Figs S2, S3**). While we did not analyze the disordered regions in detail in this study, we cannot rule out that the N-terminal structure also affects the function of RLF. It has been shown that a portion of the N-terminal region of rice RLF is similar to the N-terminal structure of human NADH cytochrome b5 oxidoreductase (Ncb5or) and adopts a helical structure when interacting with Cytb5-HBD within the same protein (Benson *et al*., 2024) How the helical structure contributes to the function of RLF is currently unknown, but the Cytb5-HBD-dependent change in the disordered region might be significant. While the N-terminal structure of each plant species is different, MpRLF rescued the LR formation phenotype of the *Arabidopsis rlf* mutant, suggesting that the disordered region may not be essential for the function of RLF. These results raise further questions about the significance of the disordered region in the RLF protein family.

In this study, we observed that the Mp*rlf^ge^* mutant exhibited inhibition of gemma cup formation and retarded thallus growth (**Figs 4, 5, S5**), which is consistent with Mp*RLF* promoter activity at the bottom of the gemma cup and the ventral side of the thallus (**Fig. 3b,c**). These results indicate that Mp*RLF* is essential for proper thallus and gemma cup formation. Several genes have been identified as essential for the formation of gemmae cups and gemma. The *GEMMA CUP-ASSOCIATED MYB1* (*GCAM1*) gene, encoding an R2R3-MYB-type transcription factor, is a master regulator of gemma cup formation, and *GCAM1* knockout lines are unable to form any gemma cup and gemma (Yasui *et al*., 2019). The *KARAPPO* gene, which encodes a ROP guanine nucleotide exchange factor (RopGEF), is also essential for gemma formation. Loss-of-function mutants of *KARAPPO* display a typical phenotype of impaired gemma formation without affecting gemma cup formation (Hiwatashi *et al*., 2019). In addition, mutants of the Mp*RKD* gene, which regulates germ cell differentiation and initiation, are defective in forming the rim of the gemma cup although gemmae still develop from the bottom of the gemma cup (Koi *et al*., 2016; Rövekamp *et al*., 2016). Compared with the above mutants, Mp*rlf^ge^* forms gemma on the thallus while proper gemma cup formation is inhibited. Interestingly, in the RNA-seq dataset, *Mp5g06550/KARAPPO* expression was unchanged in Mp*rlf^ge^*, while *Mp3g04030/*Mp*RKD* and *Mp6g04830/GCAM1* expression was decreased (**Table S3**). It should be noted that we used plants grown for 16 days, which is sufficient time for gemma cup formation in wild type plants under our growth condition. MpRLF is required for the proper expressions of *GCAM1* and Mp*RKD* during the process of gemma-cup formation. In addition to the defects in vegetative development, Mp*rlf^ge^* mutants showed aberrant development in male and female reproductive organs (**Fig. 5b**), indicating that Mp*RLF* is also essential for proper reproductive organ development. Further analyses of the expression pattern of the genes that regulate the formation of gemma cup, gemma, and reproductive organs (antheridiophore and archegoniophore) in Mp*rlf^ge^*will shed light on how Mp*RLF* regulates the development of these organs. Transcriptome analysis using the Mp*rlf^ge^* mutant also suggest that MpRLF may contribute to organ development by regulating several metabolic pathways. Regarding the *A. thaliana* Cytb5-HBD family, Cytb5-HBD proteins function primarily as an electron carrier, transferring electrons from reductases to target proteins. Specifically, CB5D provides electrons to enzymes that convert precursors of G-lignin synthesis to precursors of S-lignin synthesis, thereby regulating lignin synthesis (Gou *et al*., 2019). CB5B is involved in sphingolipid synthesis through its interaction with FAH1, which hydroxylates the C-2 position of fatty acids and provides the necessary electrons for the reaction (Nagano *et al*., 2012). These findings suggest that RLFs may also function as electron transmitters. Consistent with this, our RNA-seq analysis showed that the expression of genes involved in electron transfer and redox is altered in Mp*rlf^ge^* (**Figs 7, S7, S8**). This result suggests that MpRLF is an electron transferase that regulates the cellular redox state. Additionally, the Cytb5-HBD protein may also be an electron transferase in *M. polymorpha*, although its specific action pathway remains to be elucidated. Therefore, identifying MpRLF-interacting proteins through co-immunoprecipitation or proximity labeling using active biotin ligase (TurboID) will be valuable (Mair *et al*., 2019; Zhang *et al*., 2020).

In this study, GO enrichment and KEGG pathway analyses revealed that the variation in gene expression in Mp*rlf^ge^* is most pronounced in metabolic processes, cellular components, and molecular functions (**Fig. S7b**). Furthermore, the GO hierarchy of the top 200 downregulated genes was enriched in GO terms related to nitrate response, transport, and metabolism (**Fig. 8**, **Table S1**). Of particular interest is the fact that RLF regulates LR formation in *A. thaliana* (Ikeyama *et al*., 2010). The link between LR formation and nitrate has been extensively studied in recent years. The nitrate-responsive OBP4-XTH9 regulatory module plays a crucial role in LR development and regulates LR growth in response to changes in nitrate concentration (Xu & Cai, 2019). Furthermore, it has been demonstrated that nitrate interacts with auxin during LR formation, and that the distribution of nitrate concentration significantly affects LR development (Pélissier *et al*., 2021). We have not examined the involvement of nitrate in RLF-dependent organ development, but there may be a causative link between nitrate and abnormal organ development in Mp*rl^fge^*.

Further study of RLF orthologs in plants, including *Arabidopsis* RLF and MpRLF, will elucidate the relationship between the evolution of Cytb5-HBD and biological roles of the RLF protein family in organ growth and development. This will contribute to a deeper understanding of the evolutionarily conserved processes governing plant organ development.

## Acknowledgements

We thank Tsuyoshi Nakagawa (Shimane University, Japan) for pGWB vectors, Hirotaka Kato (Ehime University, Japan) for *E2Fp::XVE>>X-GFP* vector, and Akina Senoo and Junko Unten for technical assistance. This work was funded by grants from the Ministry of Education, Culture, Sports, Science, and Technology (MEXT), Japan: Grants-in-Aid for Scientific Research on Innovative Areas (19H05673 and 19H05670 to H Fukaki), JSPS KAKENHI grants (21H05271 and 23KK0127 to TS, 24K09497 and 24H02069 to TM, 21J40092 to YS, and 19H03247 to K Ishizaki), GteX Program Japan (JPMJGX23B0 to K Ishizaki) and JST SPRING (JPMJSP2148 to KP Iwata).

## Competing interests

None declared.

## Author contributions

KP Iwata and H Fukaki designed the study. KP Iwata performed most of the experiments. TS and TM measured the absorbance of the proteins. YS, TF, H Fukumura, and K Ishizaki supported the experiments using *M. polymorpha*. KP Iwata, TS, YS, TF, H Fukumura, YK, TM, K Ishizaki and H Fukaki discussed about the research. KP Iwata and H Fukaki wrote the manuscript, with inputs from all authors.

## Data availability

The data underlying this article are available in the article and in its online supplementary materials.

## Short legends for Supplemental figures and tables

**Fig. S1** Predicted structure of wild-type and mutant RLF proteins (**a**) Structure of *Arabidopsis thaliana* RLF as predicted by AlphaFold2 (modified from uniport; URL https://www.uniprot.org/uniprotkb/Q9LXD1/entry) showing the histidine residues H161G and H184G, which are predicted to be heme-binding sites, in the Cytb5-HBD domain. (**b**) 3D structures of wild-type and mutant RLF as predicted by colabfold (Mirdita *et al*., 2022). Their respective local Distance Difference Test (lDDT) scores are shown below the structures. Regions with high lDDT scores are considered to be very accurate predictions and are physically stable, which indicates that the predicted structure closely resembles the actual conformation. The lDDT scores of the respective C-terminal Cytb5-HBDs are above 70, indicating that this structure is very robust. (**c**) Photographs of the pellets of *E. coli* cells expressing wild-type and mutant MBP-RLF proteins; the color of wild-type MBP-RLF pellet is reddish-brown, whereas that of the mutant MBP-RLF pellet is not. The recombinant proteins obtained from these *E. coli* lines were used for the absorbance measurements in Fig. 1b.

**Fig. S2** Alignment of the amino acid sequence of the RLF orthologs Alignment of amino acid sequence of the RLF orthologs of *Arabidopsis thaliana* (At5g09680), *Oryza sativa* (Os07g0232200), *Zea mays* (LOC100272407), *Cryptomeria japonica* (XP_057826847.1), *Marchantia polymorpha* (Mp3g04390), *Physcomitrium patens* (XP_024357984.1), *Ceratopteris richardii* (KP509_01G108300), *Chlamydomonas reinhardtii* (CHLRE_13g574800v5), *Volvox reticuliferus* (Vretifemale_5060). The more highly conserved residues are colored in deep blue. Red lines indicate Cytb5-HBD, and asterisks indicate histidine residues that are predicted to coordinate to bind to heme. The alignment was generated using the MAFFT Multiple Sequence Alignment program (Katoh & Standley, 2013) and displayed using the Jalview software (Waterhouse *et al*., 2009).

**Fig. S3** The molecular phylogenetic tree of the RLF protein family The molecular phylogenetic tree based on RLF amino acid sequences. This tree was created using the maximum likelihood method (1000 bootstrap tests), the WAG (Whelan and Goldman) model (Whelan & Goldman, 2001), and MEGA11 software. The numbers above the nodes indicate bootstrap values and the numbers below the lines indicate evolutionary distances.

**Fig. S4** Expression pattern of the *Mp3g04390* (Mp*RLF*) gene in Marpolbase Expression database Marpolbase Expression, a publicly available database (URL https://marchantia.info/mbex/; (Kawamura *et al*., 2022) provides the expression data of the *Mp3g04390* (Mp*RLF*) gene. TPM stands for "Transcripts Per Million."

**Fig. S5** Mutations in Mp*RLF* inhibit gemma and thallus growth (**a**) Growth of Mp*RLF* loss-of-function mutants and wild-type gemma or thallus. Black scale bar = 1 mm, white scale bar = 5 mm. (**b**) Structure of Mp*RLFcds.* Mp*RLFcds_resistant* synonymous substitutions to prevent targeting by gRNA. Blue letters indicate substituted nucleotides. Amino acid sequence is shown below nucleotide sequence. (**c**) Sequence with mutations in Mp*rlf^ge^*were produced in the Tak-2 background. Blue letters indicate inserted bases, dash lines indicate deleted bases. We used pMpGE011 #6 for Mp*rlf^ge^*/Tak-2 in Fig. 5b.

**Fig. S6** Overexpression of MpRLF inhibits thallus growth Wild type (Tak-1) and three different lines *E2Fpro::XVE-*Mp*RLFcds-GFP/*Tak-1. Plants were grown on half-strength B5 medium for 7 days, then transferred to 10 µM estradiol (EST) containing medium or mock medium (0.1% (v/v) ethanol) and grown for another 7 days. Scale bar = 5 mm.

**Fig. S7** RNA-seq analysis between wild type and Mp*rlf^ge^* mutant (**a**) Pie chart showing the proportion of clean reads that mapped to different plant species. Each color represents a species as indicated in the legend, and the percentage reflects the proportion of total reads that mapped to that species. (**b**) GO term enriched across differentially expressed genes. (**c**) KEGG pathway analysis of all genes which expression is altered in Mp*rlf^ge^*#3.

**Fig. S8** Gene set enrichment analysis between wild type and Mp*rlf^ge^* mutant Gene set enrichment analysis (GSEA) based on molecular function (**a**) and cellular component (**b**) in the GO term. Each bubble chart shows the top 20 GO terms in log10(p_fdr_bh) resulting from GO enrichment analysis for up-regulated genes (upper) and down-regulated genes (bottom) in Mp*rlf^ge^* #3. The horizontal axis represents the number of genes counted, and the color bar indicates log10 (p p_fdr_bh).

**Table S1** Genes enriched in inorganic anion transport (GO:0015698) and chitin catabolic process (GO:0006032) in Fig. 8

**Table S2** Primers used in this study

**Table S3** Gene expression level in Mp*rlf^ge^* mutant

## Notes

### Competing Interest Statement

The authors have declared no competing interest.

## References

Benson DR, Deng B, Kashipathy MM, Lovell S, Battaile KP, Cooper A, Gao P, Fenton AW, Zhu H. 2024. The N-terminal intrinsically disordered region of Ncb5or docks with the cytochrome *b*_5_ core to form a helical motif that is of ancient origin. Proteins 92(4): 554–566.

Bowman JL, Kohchi T, Yamato KT, Jenkins J, Shu S, Ishizaki K, Yamaoka S, Nishihama R, Nakamura Y, Berger F, et al. 2017. Insights into Land Plant Evolution Garnered from the *Marchantia polymorpha* Genome. Cell 171(2): 287–304.e215.

Cheng CL, Dewdney J, Nam HG, den Boer BG, Goodman HM. 1988. A new locus (*NIA 1*) in *Arabidopsis thaliana* encoding nitrate reductase. EMBO Journal 7(11): 3309–3314.

Clough SJ, Bent AF. 1998. Floral dip: a simplified method for *Agrobacterium*-mediated transformation of *Arabidopsis thaliana*. The Plant Journal 16(6): 735–743.

Cowley AB, Altuve A, Kuchment O, Terzyan S, Zhang X, Rivera M, Benson DR. 2002. Toward Engineering the Stability and Hemin-Binding Properties of Microsomal Cytochromes *b*_5_ into Rat Outer Mitochondrial Membrane Cytochrome *b*_5_: Examining the Influence of Residues 25 and 71. Biochemistry 41(39): 11566–11581.

Ellson J, Gansner E, Koutsofios L, North SC, Woodhull G 2002. Graphviz— Open Source Graph Drawing Tools. Berlin, Heidelberg: Springer Berlin Heidelberg. 483–484.

Fernandez-Pozo N, Haas FB, Gould SB, Rensing SA. 2022. An overview of bioinformatics, genomics, and transcriptomics resources for bryophytes. Journal of Experimental Botany 73(13): 4291–4305.

Fukaki H, Nakao Y, Okushima Y, Theologis A, Tasaka M. 2005. Tissue-specific expression of stabilized SOLITARY-ROOT/IAA14 alters lateral root development in *Arabidopsis*. The Plant Journal 44(3): 382–395.

Goh T, Joi S, Mimura T, Fukaki H. 2012. The establishment of asymmetry in Arabidopsis lateral root founder cells is regulated by LBD16/ASL18 and related LBD/ASL proteins. Development 139(5): 883–893.

Gou M, Yang X, Zhao Y, Ran X, Song Y, Liu CJ. 2019. Cytochrome *b*_5_ Is an Obligate Electron Shuttle Protein for Syringe Lignin Biosynthesis in *Arabidopsis*. Plant Cell 31(6): 1344–1366.

Hederstedt L. 2012. Heme A biosynthesis. Biochimica Biophysica Acta 1817(6): 920–927.

Hiwatashi T, Goh H, Yasui Y, Koh LQ, Takami H, Kajikawa M, Kirita H, Kanazawa T, Minamino N, Togawa T. 2019. The RopGEF KARAPPO is essential for the initiation of vegetative reproduction in *Marchantia polymorpha*. Current Biology 29(20): 3525–3531.

Ikeyama Y, Tasaka M, Fukaki H. 2010. RLF, a cytochrome *b*_5_-like heme/steroid binding domain protein, controls lateral root formation independently of ARF7/19-mediated auxin signaling in *Arabidopsis thaliana*. The Plant Journal 62(5): 865–875.

Ishida S, Suzuki H, Iwaki A, Kawamura S, Yamaoka S, Kojima M, Takebayashi Y, Yamaguchi K, Shigenobu S, Sakakibara H, et al. 2022. Diminished Auxin Signaling Triggers Cellular Reprogramming by Inducing a Regeneration Factor in the Liverwort *Marchantia polymorpha*. Plant and Cell Physiology 63(3): 384–400.

Ishizaki K, Chiyoda S, Yamato KT, Kohchi T. 2008. *Agrobacterium*-Mediated Transformation of the Haploid Liverwort *Marchantia polymorpha* L., an Emerging Model for Plant Biology. Plant and Cell Physiology 49(7): 1084–1091.

Ishizaki K, Nishihama R, Ueda M, Inoue K, Ishida S, Nishimura Y, Shikanai T, Kohchi T. 2015. Development of Gateway Binary Vector Series with Four Different Selection Markers for the Liverwort *Marchantia polymorpha*. PLoS One 10(9): e0138876.

Ishizaki K, Nishihama R, Yamato KT, Kohchi T. 2016. Molecular Genetic Tools and Techniques for *Marchantia polymorpha* Research. Plant and Cell Physiology 57(2): 262–270.

Katoh K, Standley DM. 2013. MAFFT multiple sequence alignment software version 7: improvements in performance and usability. Molecular Biology Evolution 30(4): 772–780.

Kawamura S, Romani F, Yagura M, Mochizuki T, Sakamoto M, Yamaoka S, Nishihama R, Nakamura Y, Yamato KT, Bowman JL, et al. 2022. MarpolBase Expression: A Web-Based, Comprehensive Platform for Visualization and Analysis of Transcriptomes in the Liverwort *Marchantia polymorpha*. Plant and Cell Physiology 63(11): 1745–1755.

Kerfeld CA, Krogmann DW. 1998. Photosynthetic cytochromes c in cyanobacteria, algae, and plants. Annu Review of Plant Physiology and Plant Molecular Biology 49(1): 397–425.

Kim D, Langmead B, Salzberg SL. 2015. HISAT: a fast spliced aligner with low memory requirements. Nature Methods 12(4): 357–360.

Klopfenstein DV, Zhang L, Pedersen BS, Ramírez F, Warwick Vesztrocy A, Naldi A, Mungall CJ, Yunes JM, Botvinnik O, Weigel M, et al. 2018. GOATOOLS: A Python library for Gene Ontology analyses. Science Reports 8(1): 10872.

Koi S, Hisanaga T, Sato K, Shimamura M, Yamato KT, Ishizaki K, Kohchi T, Nakajima K. 2016. An Evolutionarily Conserved Plant RKD Factor Controls Germ Cell Differentiation. Current Biology 26(13): 1775–1781.

Kubota A, Ishizaki K, Hosaka M, Kohchi T. 2013. Efficient *Agrobacterium*-mediated transformation of the liverwort *Marchantia polymorpha* using regenerating thalli. Bioscience Biotechnology, and Biochemistry 77(1): 167–172.

Langmead B, Salzberg SL. 2012. Fast gapped-read alignment with Bowtie 2. Nature Methods 9(4): 357–359.

Layer G, Reichelt J, Jahn D, Heinz DW. 2010. Structure and function of enzymes in heme biosynthesis. Protein Science 19(6): 1137–1161.

Li B, Dewey CN. 2011. RSEM: accurate transcript quantification from RNA-Seq data with or without a reference genome. BMC Bioinformatics 12: 323.

Li T, Bonkovsky HL, Guo JT. 2011. Structural analysis of heme proteins: implications for design and prediction. BMC Structural Biology 11: 13.

Maggio C, Barbante A, Ferro F, Frigerio L, Pedrazzini E. 2007. Intracellular sorting of the tail-anchored protein cytochrome *b*_5_ in plants: a comparative study using different isoforms from rabbit and *Arabidopsis*. Journal of Experimental Botany 58(6): 1365–1379.

Mair A, Xu S-L, Branon TC, Ting AY, Bergmann DC. 2019. Proximity labeling of protein complexes and cell-type-specific organellar proteomes in *Arabidopsis* enabled by TurboID. Elife 8: e47864.

Mirdita M, Schütze K, Moriwaki Y, Heo L, Ovchinnikov S, Steinegger M. 2022. ColabFold: making protein folding accessible to all. Nature Methods 19(6): 679–682.

Montgomery SA, Tanizawa Y, Galik B, Wang N, Ito T, Mochizuki T, Akimcheva S, Bowman JL, Cognat V, Maréchal-Drouard L, et al. 2020. Chromatin Organization in Early Land Plants Reveals an Ancestral Association between H3K27me3, Transposons, and Constitutive Heterochromatin. Current Biology 30(4): 573–588.e577.

Nagano M, Takahara K, Fujimoto M, Tsutsumi N, Uchimiya H, Kawai-Yamada M. 2012. Arabidopsis sphingolipid fatty acid 2-hydroxylases (AtFAH1 and AtFAH2) are functionally differentiated in fatty acid 2-hydroxylation and stress responses. Plant Physiology 159(3): 1138–1148.

Okushima Y, Fukaki H, Onoda M, Theologis A, Tasaka M. 2007. ARF7 and ARF19 regulate lateral root formation via direct activation of *LBD/ASL* genes in *Arabidopsis*. Plant Cell 19(1): 118–130.

Pertea M, Pertea GM, Antonescu CM, Chang TC, Mendell JT, Salzberg SL. 2015. StringTie enables improved reconstruction of a transcriptome from RNA-seq reads. Nature Biotechnology 33(3): 290–295.

Pélissier PM, Motte H, Beeckman T. 2021. Lateral root formation and nutrients: nitrogen in the spotlight. Plant Physiology 187(3): 1104–1116.

Reedy CJ, Gibney BR. 2004. Heme Protein Assemblies. Chemical Reviews 104(2): 617–650.

Roper JM, Smith AG. 1997. Molecular localisation of ferrochelatase in higher plant chloroplasts. European Journal of Biochemistry 246(1): 32–37.

Rövekamp M, Bowman JL, Grossniklaus U. 2016. *Marchantia* MpRKD Regulates the Gametophyte-Sporophyte Transition by Keeping Egg Cells Quiescent in the Absence of Fertilization. Current Biology 26(13): 1782–1789.

Schenkman JB, Jansson I. 2003. The many roles of cytochrome *b*_5_. Pharmacology & Therapeutics 97(2): 139–152.

Schindelin J, Arganda-Carreras I, Frise E, Kaynig V, Longair M, Pietzsch T, Preibisch S, Rueden C, Saalfeld S, Schmid B, et al. 2012. Fiji: an open-source platform for biological-image analysis. Nature Methods 9(7): 676–682.

Shanmugabalaji V, Grimm B, Kessler F. 2020. Characterization of a Plastoglobule-Localized SOUL4 Heme-Binding Protein in *Arabidopsis thaliana*. Frontiers in Plant Science 11.

Shimizu T, Yasuda R, Mukai Y, Tanoue R, Shimada T, Imamura S, Tanaka K, Watanabe S, Masuda T. 2020. Proteomic analysis of haem-binding protein from Arabidopsis thaliana and Cyanidioschyzon merolae. Philosophical Transactions of the Royal Society B 375(1801): 2019.0488.

Smith MA, Napier JA, Stymne S, Tatham AS, Shewry PR, Stobart AK. 1994. Expression of a biologically active plant cytochrome *b*_5_ in *Escherichia coli*. Biochemical Journal 303 (Pt 1)(Pt 1): 73–79.

Sperling P, Zähringer U, Heinz E. 1998. A sphingolipid desaturase from higher plants. Identification of a new cytochrome b5 fusion protein. Journal of Biological Chemistry 273(44): 28590–28596.

Sugano SS, Nishihama R. 2018. CRISPR/Cas9-Based Genome Editing of Transcription Factor Genes in *Marchantia polymorpha*. Methods in Molecular Biology 1830: 109–126.

Sugano SS, Nishihama R, Shirakawa M, Takagi J, Matsuda Y, Ishida S, Shimada T, Hara-Nishimura I, Osakabe K, Kohchi T. 2018. Efficient CRISPR/Cas9-based genome editing and its application to conditional genetic analysis in *Marchantia polymorpha*. PLoS One 13(10): e0205117.

Takahashi S, Ogawa T, Inoue K, Masuda T. 2008. Characterization of cytosolic tetrapyrrole-binding proteins in *Arabidopsis thaliana*. Photochemical & Photobiological Sciences 7(10): 1216–1224.

Tsuboyama S, Nonaka S, Ezura H, Kodama Y. 2018. Improved G-AgarTrap: A highly efficient transformation method for intact gemmalings of the liverwort *Marchantia polymorpha*. Scientific Reports 8(1): 10800.

Wahl B, Reichmann D, Niks D, Krompholz N, Havemeyer A, Clement B, Messerschmidt T, Rothkegel M, Biester H, Hille R, et al. 2010. Biochemical and spectroscopic characterization of the human mitochondrial amidoxime reducing components hmARC-1 and hmARC-2 suggests the existence of a new molybdenum enzyme family in eukaryotes. Journal of Biological Chemistry 285(48): 37847–37859.

Waterhouse AM, Procter JB, Martin DM, Clamp M, Barton GJ. 2009. Jalview Version 2--a multiple sequence alignment editor and analysis workbench. Bioinformatics 25(9): 1189–1191.

Whelan S, Goldman N. 2001. A general empirical model of protein evolution derived from multiple protein families using a maximum-likelihood approach. Molecular Biology and Evolution 18(5): 691–699.

Wilkinson JQ, Crawford NM. 1991. Identification of the *Arabidopsis CHL3* gene as the nitrate reductase structural gene *NIA2*. Plant Cell 3(5): 461–471.

Xu P, Cai W. 2019. Nitrate-responsive OBP4-XTH9 regulatory module controls lateral root development in *Arabidopsis thaliana*. PLoS Genetics 15(10): e1008465.

Yang XH, Xu ZH, Xue HW. 2005. Arabidopsis membrane steroid binding protein 1 is involved in inhibition of cell elongation. Plant Cell 17(1): 116–131.

Yasui Y, Tsukamoto S, Sugaya T, Nishihama R, Wang Q, Kato H, Yamato KT, Fukaki H, Mimura T, Kubo H. 2019. GEMMA CUP-ASSOCIATED MYB1, an ortholog of axillary meristem regulators, is essential in vegetative reproduction in *Marchantia polymorpha*. Current Biology 29(23): 3987–3995.

Zhang Y, Li Y, Yang X, Wen Z, Nagalakshmi U, Dinesh-Kumar SP. 2020. TurboID-based proximity labeling for in planta identification of protein-protein interaction networks. Journal of Visualized Experiment (159): e60728.

