## Supplemental Figures, and Tables 1 & 2 for "Evolutionary conserved RLF, a plant cytochrome *b*_5_-like heme-binding protein, is essential for organ development in *Marchantia polymorpha*"

### Fig. S1

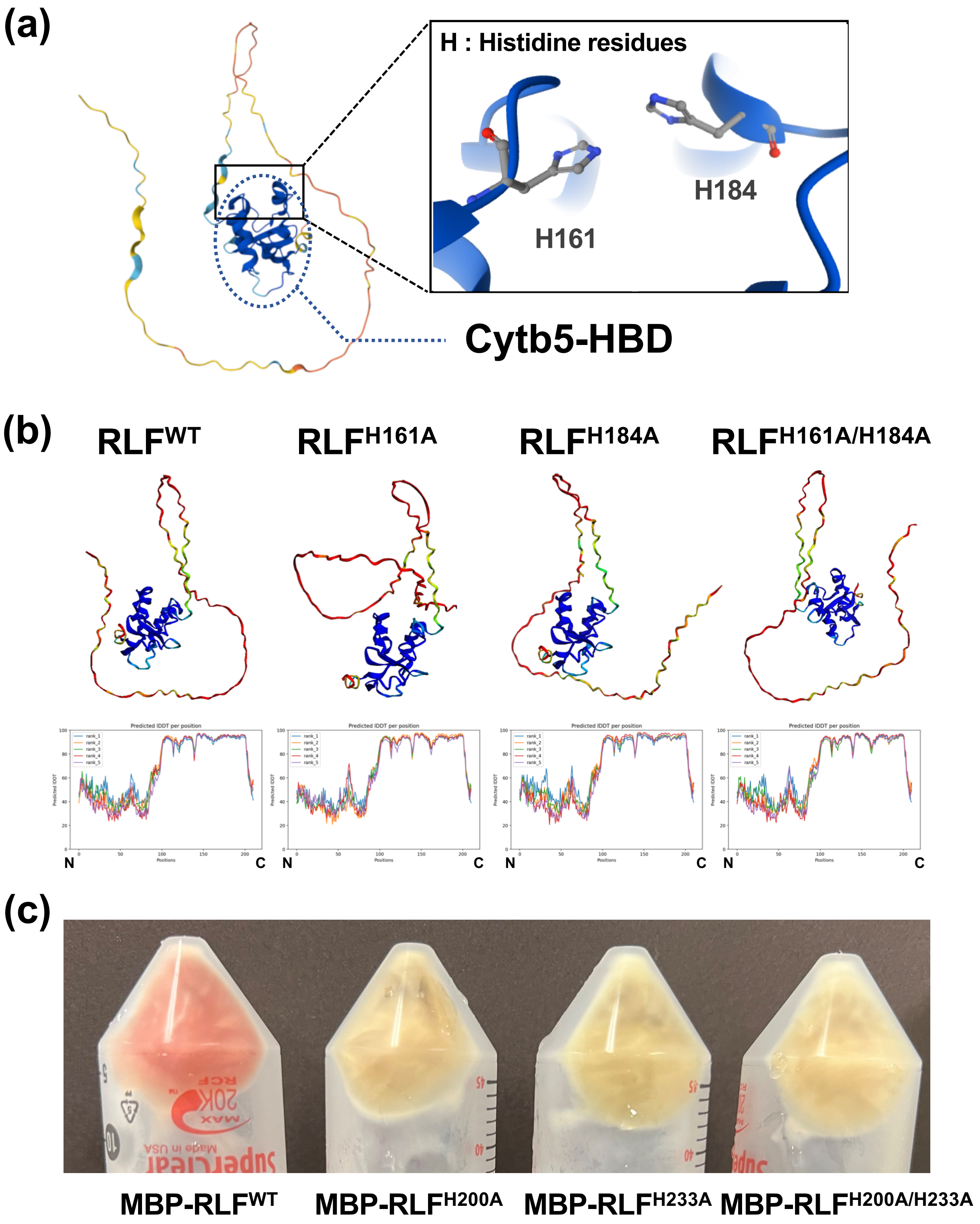

(a)
H : Histidine residues
H161
H184
Cytb5-HBD
(b)
RLF$^{WT}$
RLF$^{H161A}$
RLF$^{H184A}$
RLF$^{H161A/H184A}$
N
C
N
C
N
C
N
C
(c)
MBP-RLF$^{WT}$
MBP-RLF$^{H200A}$
MBP-RLF$^{H233A}$
MBP-RLF$^{H200A/H233A}$

### Fig. S2

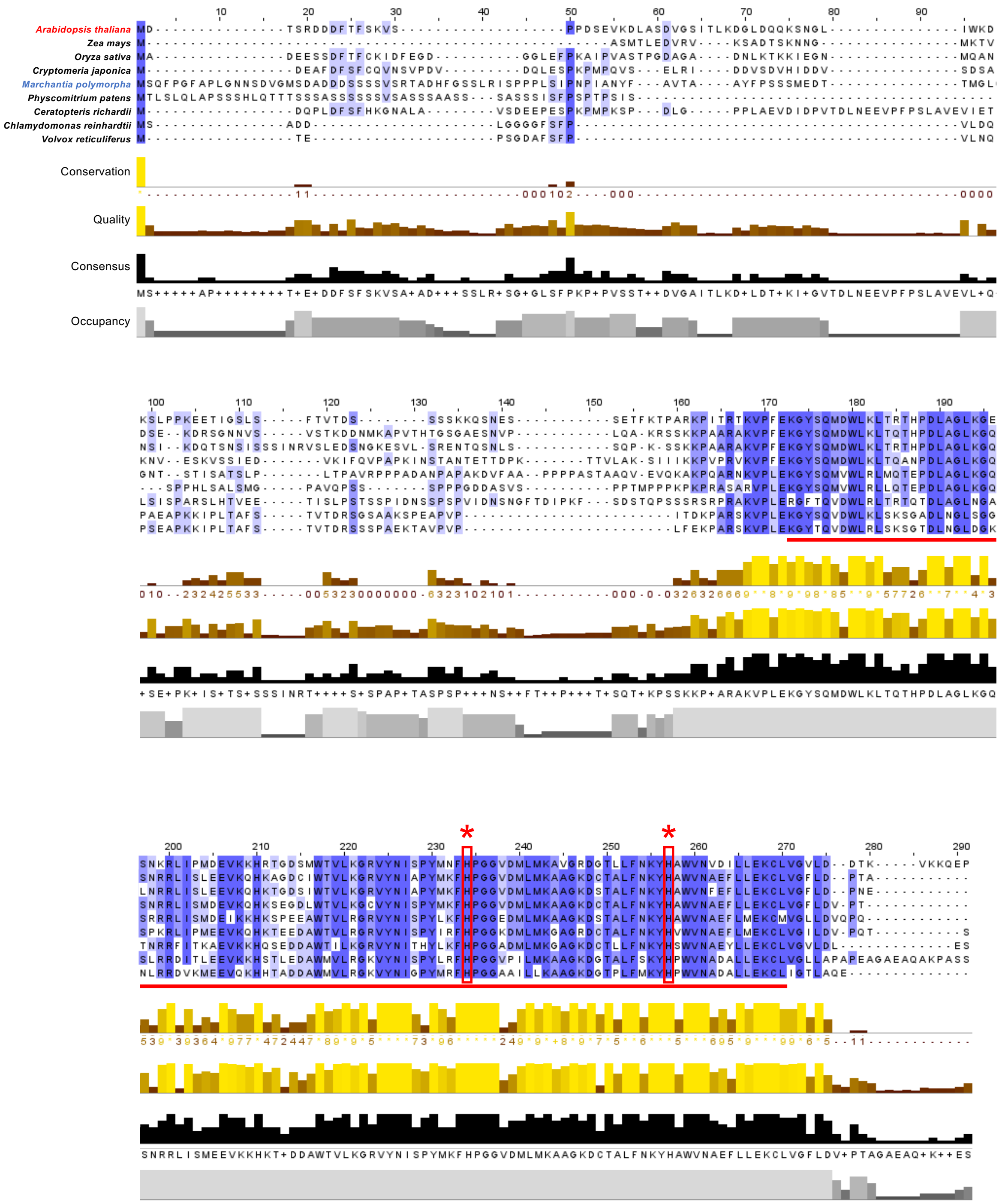

Arabidopsis thaliana
Zea mays
Oryza sativa
Cryptomeria japonica
Marchantia polymorpha
Physcomitrium patens
Ceratopteris richardii
Chlamydomonas reinhardtii
Volvox reticuliferus
Conservation
Quality
Consensus
Occupancy

### Fig. S3

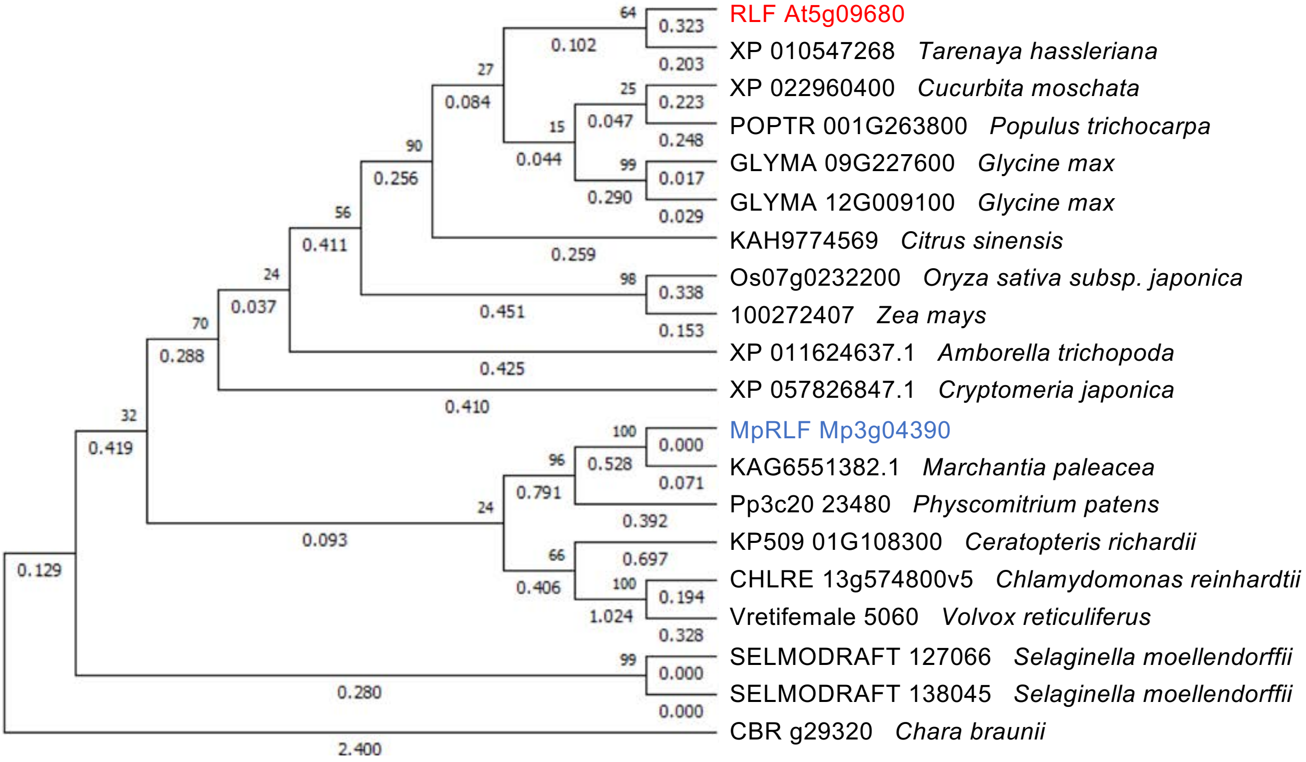
RLF At5g09680
XP 010547268 Tarenaya hassleriana
XP 022960400 Cucurbita moschata
POPTR 001G263800 Populus trichocarpa
GLYMA 09G227600 Glycine max
GLYMA 12G009100 Glycine max
KAH9774569 Citrus sinensis
Os07g0232200 Oryza sativa subsp. japonica
100272407 Zea mays
XP 011624637.1 Amborella trichopoda
XP 057826847.1 Cryptomeria japonica
MpRLF Mp3g04390
KAG6551382.1 Marchantia paleacea
Pp3c20 23480 Physcomitrium patens
KP509 01G108300 Ceratopteris richardii
CHLRE 13g574800v5 Chlamydomonas reinhardtii
Vretifemale 5060 Volvox reticuliferus
SELMODRAFT 127066 Selaginella moellendorffii
SELMODRAFT 138045 Selaginella moellendorffii
CBR g29320 Chara braunii

**Fig. S4**

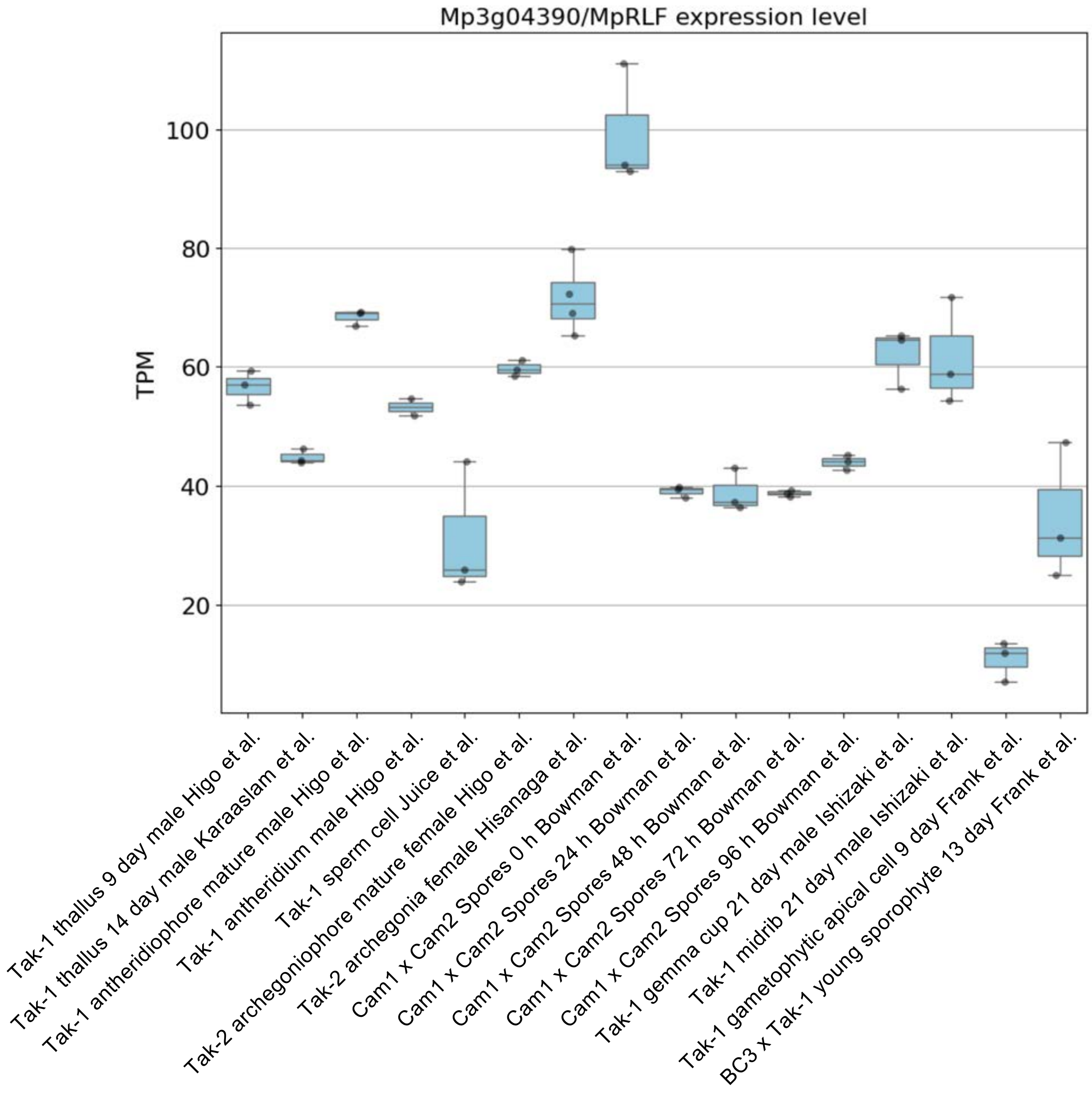

Mp3g04390/MpRLF expression level
TPM
100
80
60
40
20
Tak-1 thallus 9 day male Higo et al.
Tak-1 thallus 14 day male Karaaslam et al.
Tak-1 antheridiophore mature male Higo et al.
Tak-1 antheridium male Higo et al.
Tak-1 sperm cell Juice et al.
Tak-2 archegoniophore mature female Higo et al.
Tak-2 archegonia female Hisanaga et al.
Cam1 x Cam2 Spores 0 h Bowman et al.
Cam1 x Cam2 Spores 24 h Bowman et al.
Cam1 x Cam2 Spores 48 h Bowman et al.
Cam1 x Cam2 Spores 72 h Bowman et al.
Cam1 x Cam2 Spores 96 h Bowman et al.
Tak-1 gemma cup 21 day male Ishizaki et al.
Tak-1 midrib 21 day male Ishizaki et al.
Tak-1 gametophytic apical cell 9 day Frank et al.
BC3 x Tak-1 young sporophyte 13 day Frank et al.

### Fig. S5

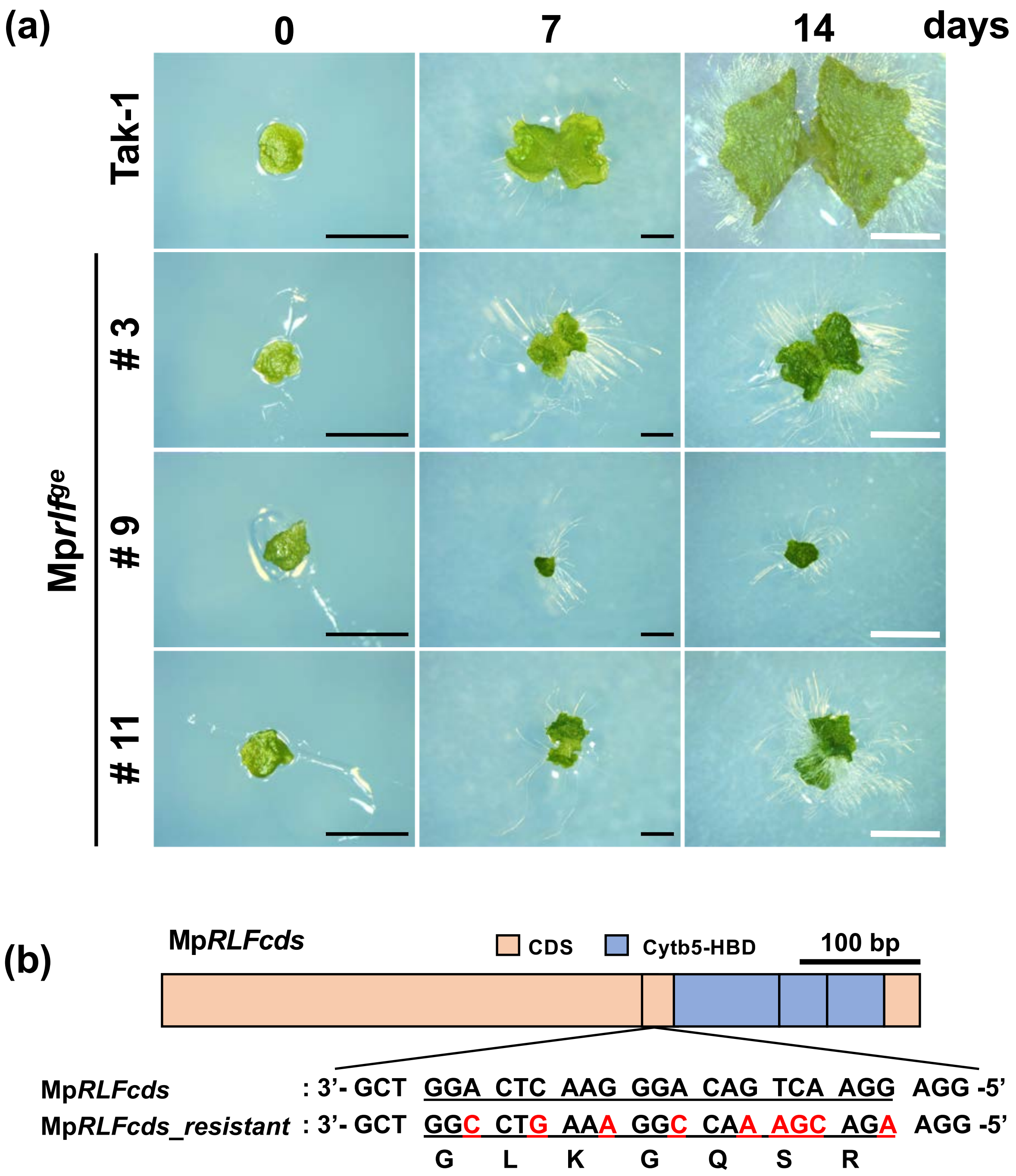


(c)

```
Tak-1       :  5'-TTCAGGACTCAAGGGACAGTCAAGGAGGAGG-3'
Tak-2       :  5'-TTCAGGACTCAAGGGACAGTCAAGGAGGAGG-3'
pMpGE010 #6  :  5'-TTCAGGACTCAAGGGACAGGTACTCAAGGAGGAGG-3'
pMpGE010 #13 :  5'-TTCAGGACTCAAGGGAC------------TGG-3'
pMpGE011 #2  :  5'-TTCAGGACTCAAGGGACA-TCAAGGAGGAGG-3'
pMpGE011 #5  :  5'-TTCAGGACTCAAGGGA-----------GGAGG-3'
pMpGE011 #6  :  5'-TTCAGGACTCAAGGGACA-TCAAGGAGGAGG-3'
pMpGE011 #9  :  5'-TTCAGGACTCAAGGGACA-TCAAGGAGGAGG-3'
pMpGE011 #10 :  5'-TTCAGGACTCAAGGGACAGGTCAAGGAGGAGG-3'
pMpGE011 #12 :  5'-TTCAGGACTCAAGGGACA-TCAAGGAGGAGG-3'
pMpGE011 #13 :  5'-TTCAGGACTCAAGGGACAGGTCAAGGAGGAGG-3'
pMpGE011 #15 :  5'-TTCAGGACTCAAGG------------AGGAGG-3'
```

Fig. S6

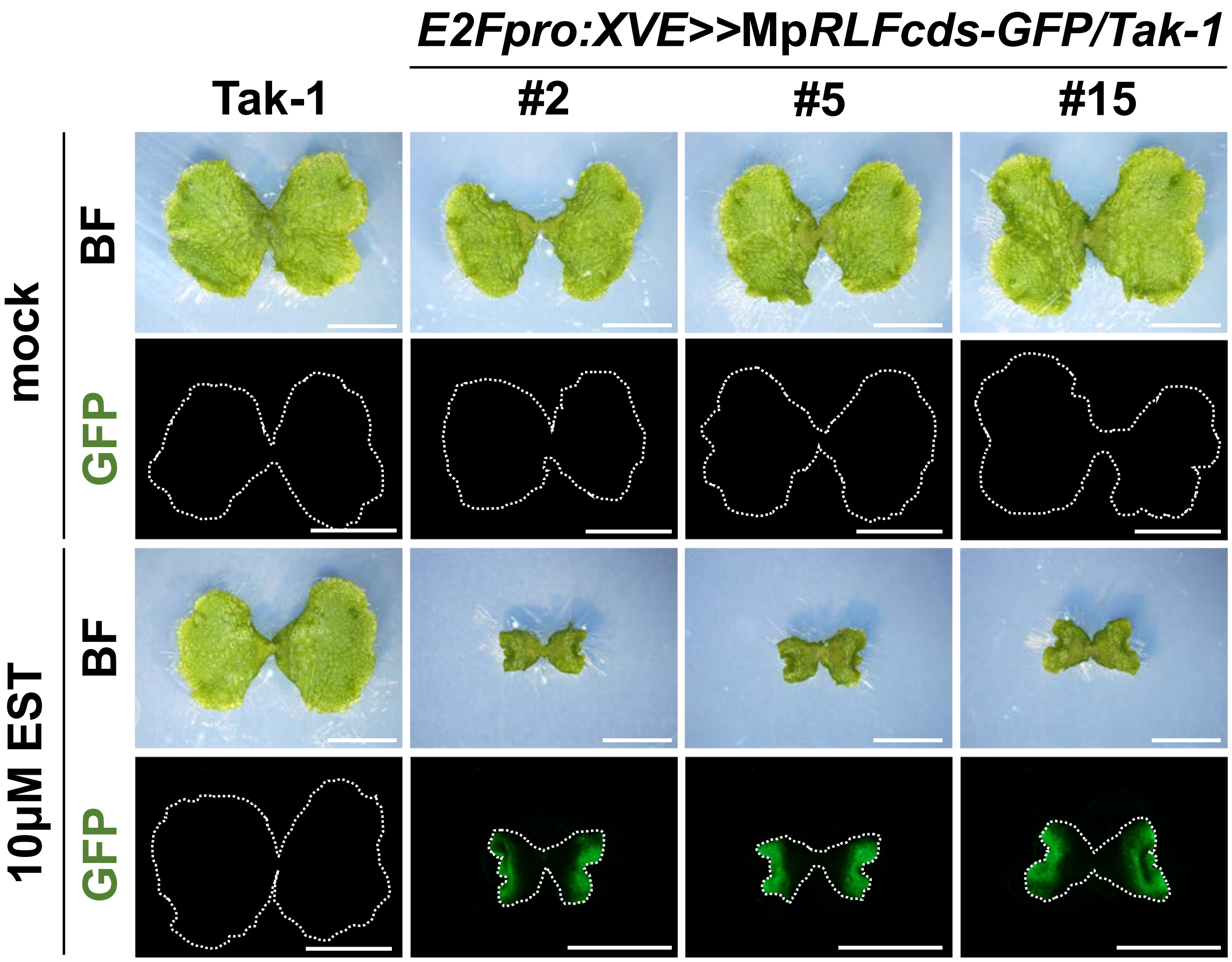

E2Fpro:XVE>>MpRLFcds-GFP/Tak-1
Tak-1
#2
#5
#15
mock
BF
GFP
10μM EST
BF
GFP

#### Fig. S7

(a)

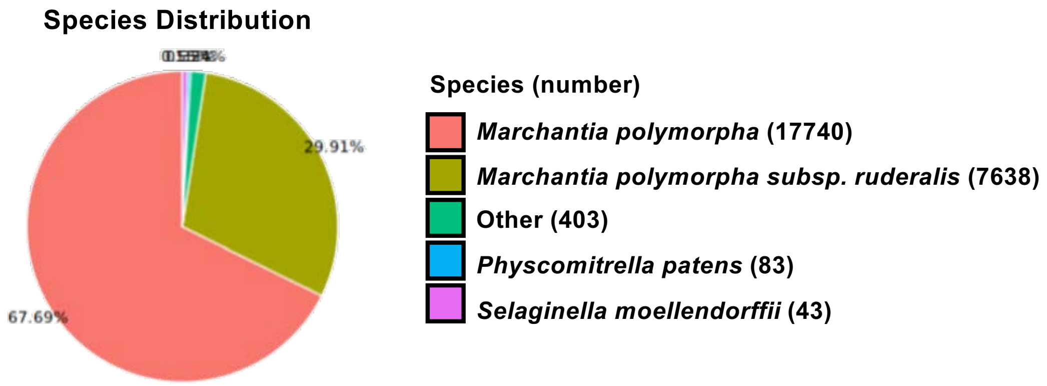

Species Distribution
29.91%
67.69%
Species (number)
Marchantia polymorpha (17740)
Marchantia polymorpha subsp. ruderalis (7638)
Other (403)
Physcomitrella patens (83)
Selaginella moellendorffii (43)


(b)

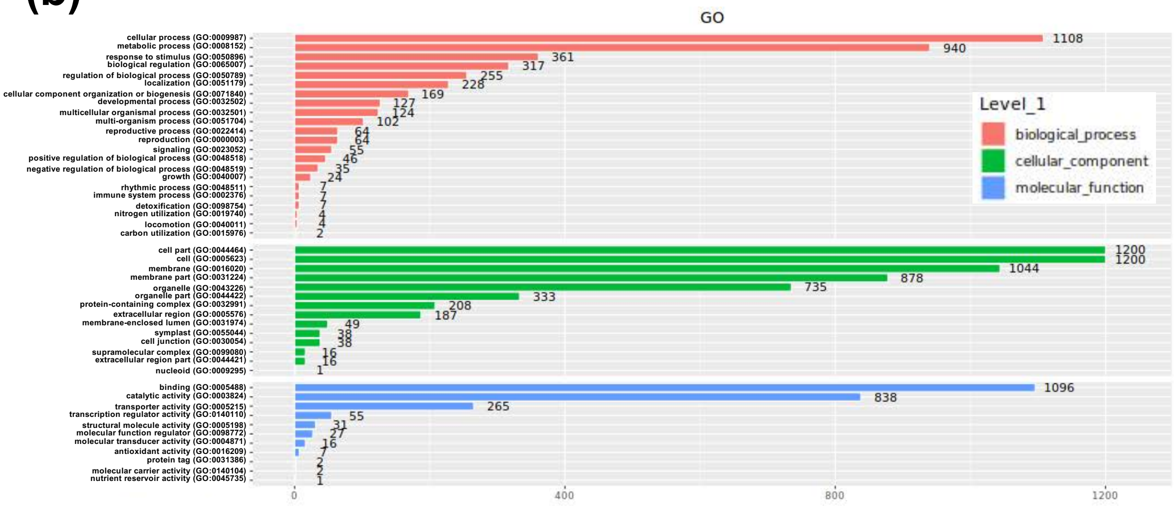

GO
cellular process (GO:0009987) 1108
metabolic process (GO:0008152) 940
response to stimulus (GO:0050896) 361
biological regulation (GO:0065007) 317
regulation of biological process (GO:0050789) 255
localization (GO:0051179) 228
cellular component organization or biogenesis (GO:0071840) 169
developmental process (GO:0032502) 127
multicellular organismal process (GO:0032501) 124
multi-organism process (GO:0051704) 102
reproductive process (GO:0022414) 64
reproduction (GO:0000003) 64
signaling (GO:0023052) 55
positive regulation of biological process (GO:0048518) 46
negative regulation of biological process (GO:0048519) 35
growth (GO:0040007) 24
rhythmic process (GO:0048511) 7
immune system process (GO:0002376) 7
detoxification (GO:0098754) 7
nitrogen utilization (GO:0019740) 4
locomotion (GO:0040011) 4
carbon utilization (GO:0015976) 2
cell part (GO:0044464) 1200
cell (GO:0005623) 1200
membrane (GO:0016020) 1044
membrane part (GO:0031224) 878
organelle (GO:0043226) 735
organelle part (GO:0044422) 333
protein-containing complex (GO:0032991) 208
extracellular region (GO:0005576) 187
membrane-enclosed lumen (GO:0031974) 49
symplast (GO:0055044) 38
cell junction (GO:0030054) 38
supramolecular complex (GO:0099080) 16
extracellular region part (GO:0044421) 16
nucleoid (GO:0009295) 1
binding (GO:0005488) 1096
catalytic activity (GO:0003824) 838
transporter activity (GO:0005215) 265
transcription regulator activity (GO:0140110) 55
structural molecule activity (GO:0005198) 31
molecular function regulator (GO:0098772) 27
molecular transducer activity (GO:0004871) 16
antioxidant activity (GO:0016209) 7
protein tag (GO:0031386) 2
molecular carrier activity (GO:0140104) 2
nutrient reservoir activity (GO:0045735) 1
0
400
800
1200
Level_1
biological_process
cellular_component
molecular_function


(c)

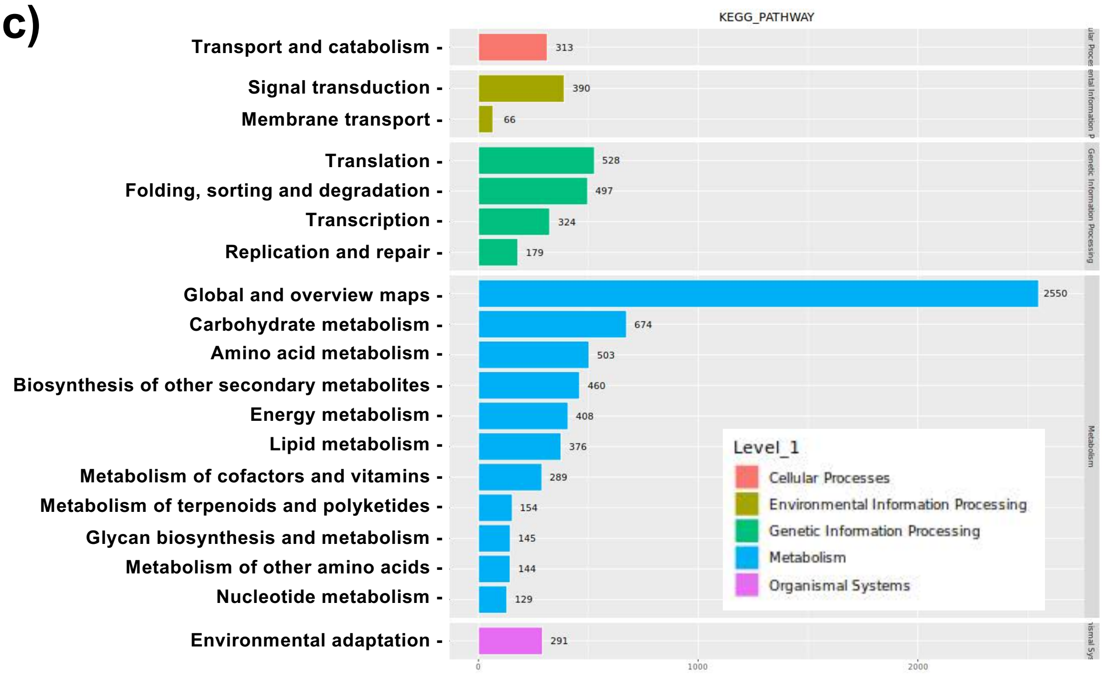

KEGG_PATHWAY
Transport and catabolism 313
Signal transduction 390
Membrane transport 66
Translation 528
Folding, sorting and degradation 497
Transcription 324
Replication and repair 179
Global and overview maps 2550
Carbohydrate metabolism 674
Amino acid metabolism 503
Biosynthesis of other secondary metabolites 460
Energy metabolism 408
Lipid metabolism 376
Metabolism of cofactors and vitamins 289
Metabolism of terpenoids and polyketides 154
Glycan biosynthesis and metabolism 145
Metabolism of other amino acids 144
Nucleotide metabolism 129
Environmental adaptation 291
0
1000
2000
Genetic Information Processing
Metabolism
Level_1
Cellular Processes
Environmental Information Processing
Genetic Information Processing
Metabolism
Organismal Systems

### Fig. S8

#### (a) Molecular Function

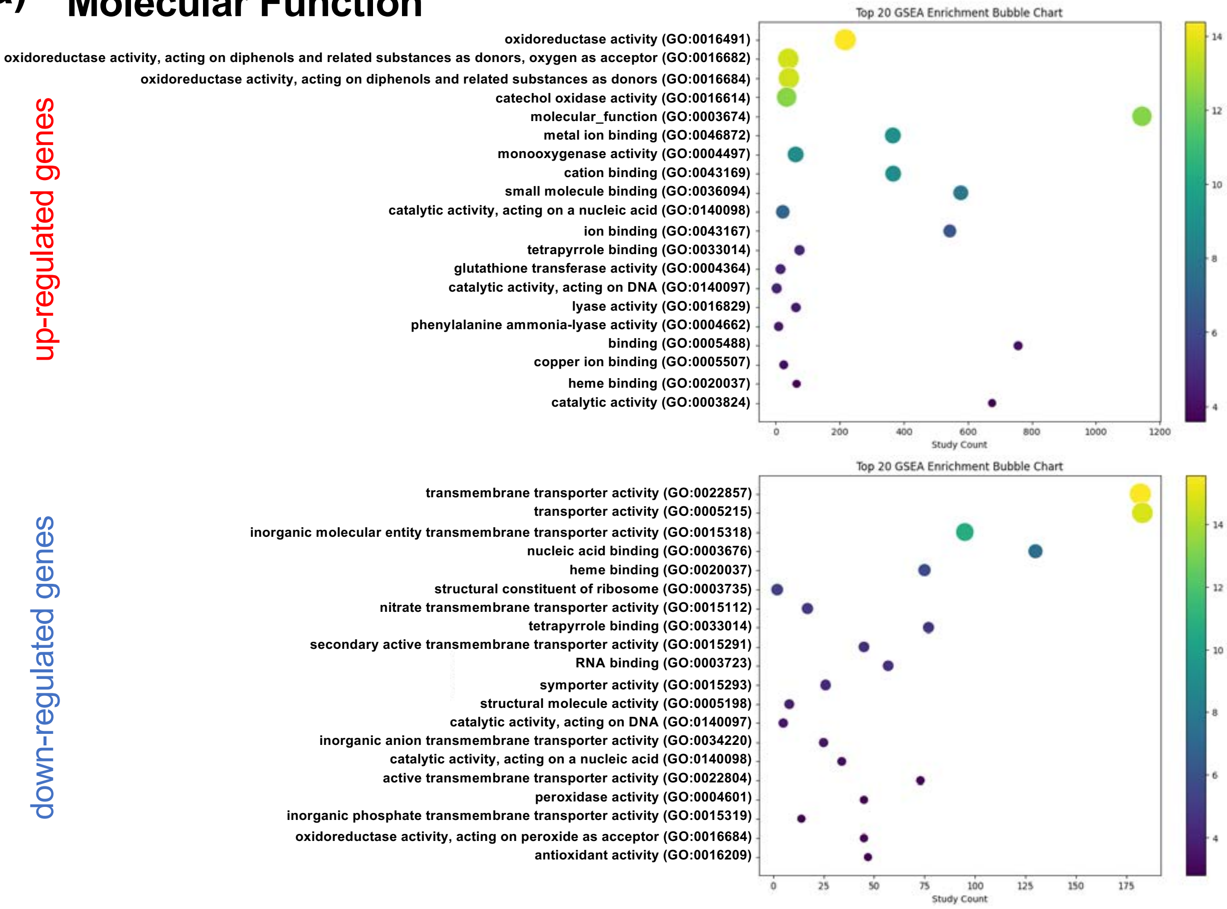


#### (b) Cellular Component

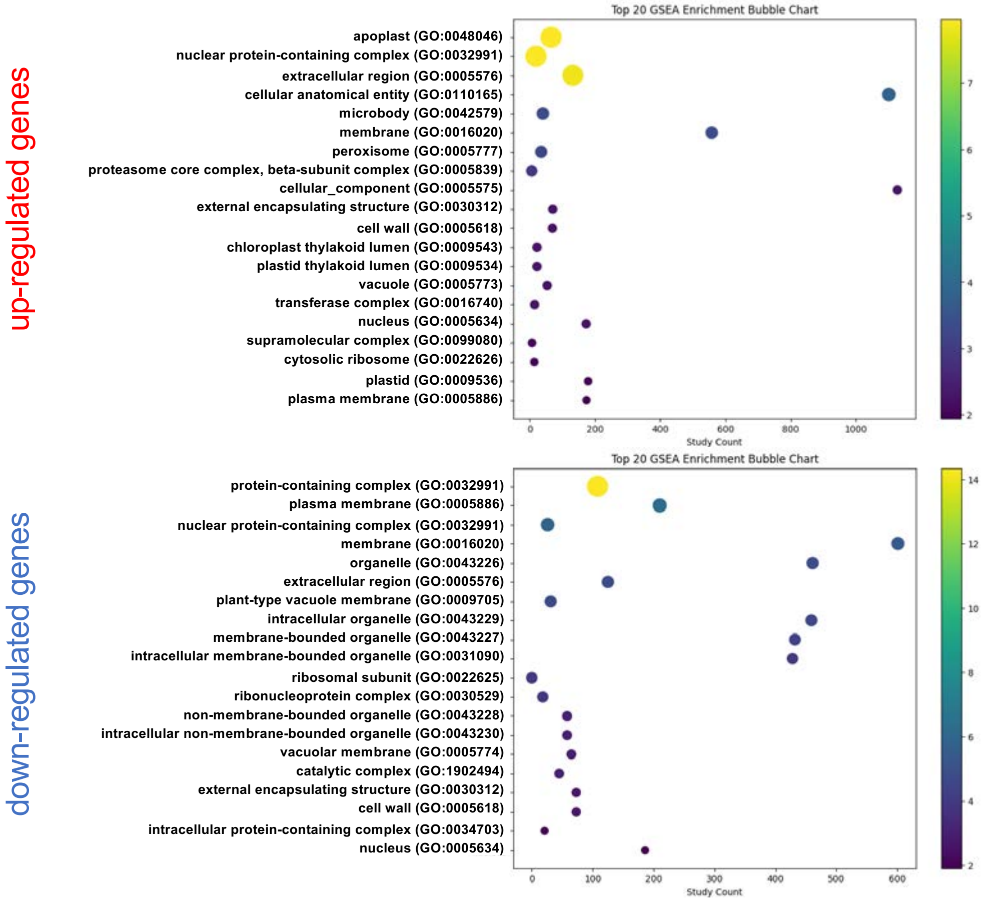

### Table S1

**Table S1 Genes enriched in inorganic anion transport (GO:0015698) and chitin catabolic process (GO:0006032) in Fig. 8**

| GO:0015698 | inorganic anion transport |
|---|---|
| *Mp1g17070* | |
| *Mp2g05910* | |
| *Mp3g14570* | |
| *Mp4g16610*/Mp*PTB5* | Phosphate Transporter |
| *Mp4g22680* | High-affinity Nitrate Transporter 2.3 |
| *Mp4g22700* | High Affinity Nitrate Transporter 2.4 |
| *Mp5g07750* | High Affinity Nitrate Transporter 2.4 |
| *Mp5g10710* | High Affinity Nitrate Transporter 2.4 |
| *Mp8g04800* | High Affinity Nitrate Transporter 2.4 |

| GO:0006032 | chitin catabolic process |
|---|---|
| *Mp1g22460*/Mp*GH18.1* | Cell wall related protein |
| *Mp4g01780*/Mp*GH19.4* | Cell wall related protein |
| *Mp4g20440*/Mp*GH19.6* | Cell wall related protein |
| *Mp4g20450*/Mp*GH19.7* | Cell wall related protein |
| *Mp4g20470*/Mp*GH19.8* | Cell wall related protein |
| *Mp5g02510*/Mp*GH19.13* | Cell wall related protein |

### Table S2

#### Table S2 Primers used in this study

| Name | Primer sequence (5'-3') |
| --- | --- |
| CACC-RLF-F | CACCATGGATACTAGTAGAGATGAT |
| RLF-XhoI-R | TCGAGGGGTTCTTGCTTCTTCACTTT |
| IF-pGWB501RLFpro-F | GGCCAGTGCCAAGCTTGACAAGAGAGGACAATTTGG |
| IF-pGWB501RLFpro-R | GCAGGCATGCAAGCTCTTGAGCTTAATCTGCACGC |
| H161G-F | AATTTTGGTCCTGGAGGTGTCGATATG |
| H161G-R | TCCAGGACCAAAATTCATATAAGGCGA |
| H184G-F | AAATACGGTGCCTGGGTCAATGTTGAT |
| H184G-R | CCAGGCACCGTATTTGTTGAATAAGAG |
| H161A-F | AATTTTGCTCCTGGAGGTGTCGATATG |
| H161A-R | TCCAGGAGCAAAATTCATATAAGGCGAGC |
| H184A-F | AAATACGCTGCCTGGGTCAATGTTGAT |
| H184A-R | CCAGGCAGCGTATTTGTTGAATAAGAG |
| MpRLFpro-CACC-F | CACCACTCGGTGGTATTCAAGCACTCA |
| MpRLFpro-R | CATGGTCCACGTCTGTTACGC |
| MpRLF-CRISPR-F | CTCGGGACTCAAGGGACAGTCA |
| MpRLF-CRISPR-R | AAACTGACTGTCCCTTGAGTCC |
| MpRLFcds-CACC-F | CACCATGTCTCAGTTTCCTGGTTTTGCG |
| IF-MpRLFpro-F | GGCCAGTGCCAAGCTACTCGGTGGTATTCAAGCACTCA |
| IF-MpRLFpro-R | CTCTAGACCCAAGCTGGTCCACGTCTGTTACGC |
| MpRLFcds(-stop)-R | TTGTGGTTGGACGTCCAGC |
| MpRLFcds(+stop)-R | TCATTGTGGTTGGACGTCCA |
| IF-MpRLF(com)-F | GGCCTGAAAGGCCAAAGCAGAAGGAGGTTGATCAGTATG |
| IF-MpRLF(com)-R | TCTGCTTTGGCCTTTCAGGCCAGCAAGATCAGGTTCAGT |
| RLFcds-CACC-F | CACCATGGATACTAGTAGAGATGAT |
| XhoI-RLF-R | CTCGAGGGGTTCTTGCTTCTTCACTTT |
| IF-RLFpro-F | GGCCAGTGCCAAGCTTGACAAGAGAGGACAATTTGG |
| IF-RLFpro-R | GCAGGCATGCAAGCTCTTGAGCTTAATCTGCACGCA |
| XhoI-GFP-F | GGCCTCGAGGGAGGCGGTGGAGGCATGGTG |
| SalI-GFP(stop)-R | GGCGTCGACCTGCAGGCGGCCGCG |

**Fig. S1** Predicted structure of wild-type and mutant RLF proteins
